# Ceramide-Rich Microdomains Facilitate Nuclear Envelope Budding during the Biogenesis of LTB_4_-containing Exosomes

**DOI:** 10.1101/2022.02.09.479761

**Authors:** Subhash B. Arya, Song Chen, Fatima Javed, Carole A. Parent

**Author notes:** Corresponding Author: Life Sciences Institute, University of Michigan, 210 Washtenaw Ave, Office 4437, Ann Arbor, MI 48109-2216.

## Abstract

Neutrophils migrating towards chemoattractant gradients amplify their recruitment range by releasing the secondary chemoattractant leukotriene B_4_ (LTB_4_)^1,2^. We previously demonstrated that LTB_4_ and its synthesizing enzymes, the 5-lipoxygenase (5-LO), 5-LO activating protein (FLAP), and leukotriene A_4_ hydrolase (LTA_4_H), are packaged and released in exosomes^3^. We now report that the biogenesis of the LTB_4_-containing exosomes is initiated at the nuclear envelope (NE) of activated neutrophils. We show that the neutral sphingomyelinase 1 (nSMase1)-mediated generation of ceramide enriched lipid-ordered microdomains initiates the clustering of the LTB_4_-synthesizing enzymes on the NE. We isolated and analyzed exosomes from activated neutrophils and established that the FLAP/5-LO-positive exosome population is distinct from that of the CD63-positive exosome population. Furthermore, we observed a strong co-localization between ALIX and FLAP at the periphery of nuclei and within cytosolic vesicles. We propose that the initiation of NE curvature and bud formation is mediated by nSMase1-dependent ceramide generation, which leads to FLAP and ALIX recruitment. Together, these observations elucidate the mechanism for LTB_4_ secretion and identify a novel pathway for exosome generation.

---

Neutrophils represent the first line of defense at sites of injury or infection^4^. Upon exposure to primary chemoattractants, e.g., formyl peptide N-Formyl-methionine-leucyl-phenylalanine (fMLF), neutrophils rapidly secrete the secondary chemoattractant leukotriene B_4_ (LTB_4_) which serves to maintain the robustness and sensitivity to primary chemoattractant signals and dramatically increase the range and persistence of migration^5^. LTB_4_-synthesis is initiated with the release of arachidonic acid (AA) through phospholipid hydrolysis mediated by the translocation of cytosolic-phospholipase A_2_ (cPLA_2_) to the nuclear envelope (NE)^6^. The NE-associated transmembrane protein, 5-lipoxygenase activating protein (FLAP), presents the released AA to 5-lipoxygenase (5-LO), which generates leukotriene A_4_ (LTA_4_). LTA_4_ is in turn quickly hydrolyzed to LTB_4_ by LTA_4_ hydrolase (LTA_4_H)^7^. It has been shown that the LTB_4_-synthesizing enzymes are present in exosomes secreted from macrophages, dendritic cells, and chemoattractant-activated neutrophils^3,8^. Exosomes are synthesized as intraluminal vesicles (ILVs) within multivesicular bodies (MVBs) and secreted upon fusion of MVBs with the plasma membrane. The sorting of cargos packaged in ILVs is mediated by the tetraspanin CD63, the endosomal sorting complex required for transport (ESCRT) complexes, or affinity towards neutral sphingomyelinase (nSMase)-dependent ceramide-rich lipid microdomains^9^. The inhibition of LTB_4_ synthesis *ex vivo*^5^ or *in vivo*^10,^11,12 or the inhibition of exosome release through neutral sphingomyelinase1 (nSMase1) knockdown^3^ results in diminished neutrophil recruitment in response to sterile injury or bacterial peptides with a concomitant decrease in recruitment range. In this context, we envision that the packaging of LTB_4_ in exosomes provides a mechanism to support intercellular signaling in harsh extracellular environments.

To gain more insight into the mechanisms that regulate the biogenesis of LTB_4_-containing exosomes, we measured FLAP and 5-LO distribution in activated human peripheral blood-derived primary polymorphonuclear neutrophils (PMNs) uniformly stimulated with fMLF. In resting cells, while FLAP clearly showed NE and reticulate (endoplasmic reticulum) localization, 5-LO mainly localized inside the nucleus. However, 15 min after fMLF addition, we readily observed the appearance of FLAP-positive NE buds that also contained 5-LO (**Supplementary Fig. 1A**). The NE origin of these buds was confirmed using the inner-nuclear membrane (INM) resident protein, Lamin B Receptor (LBR), which we observed around 5-LO-positive perinuclear as well as cytosolic vesicles (**Fig. 1A**). The temporal increase in the percentage of PMNs with LBR-positive NE buds and cytosolic vesicles upon fMLF stimulation highlights the requirement for chemotactic activation in the generation of NE-derived buds and cytosolic vesicles (**Fig. 1B**). Similar findings were observed in PMNs chemotaxing towards fMLF (**Fig. 1C&D**), where we observed that the LBR-positive vesicles present in the cytosol were consistently smaller in size, compared to the NE-associated buds in these cells (**Fig. 1C, D and Supplementary Fig. 1B**). We also observed the presence of LTA_4_H within the LBR-positive NE-buds and cytosolic vesicles (**Fig. 1E&F**), substantiating the nuclear origin of the LTB_4_-containing exosomes. NE budding and vesiculation have been reported during the formation of micronuclei in cancer cells and are characterized by the presence of the nucleoskeleton proteins, LaminB1 and or Lamin A/C, along with the DNA repair machinery in LBR-positive vesicles^13,14^. To rule out the possibility that the LBR-positive NE buds and cytosolic vesicles we observed are micronuclei, we immunostained chemotaxing PMNs with Lamin B1 as PMNs express very low levels of Lamin A/C and are devoid of measurable DNA repair^15^. We found that the LBR positive buds and cytosolic vesicles are devoid of Lamin B1 signal (**Supplementary Fig. 1C**).

**Figure 1.**
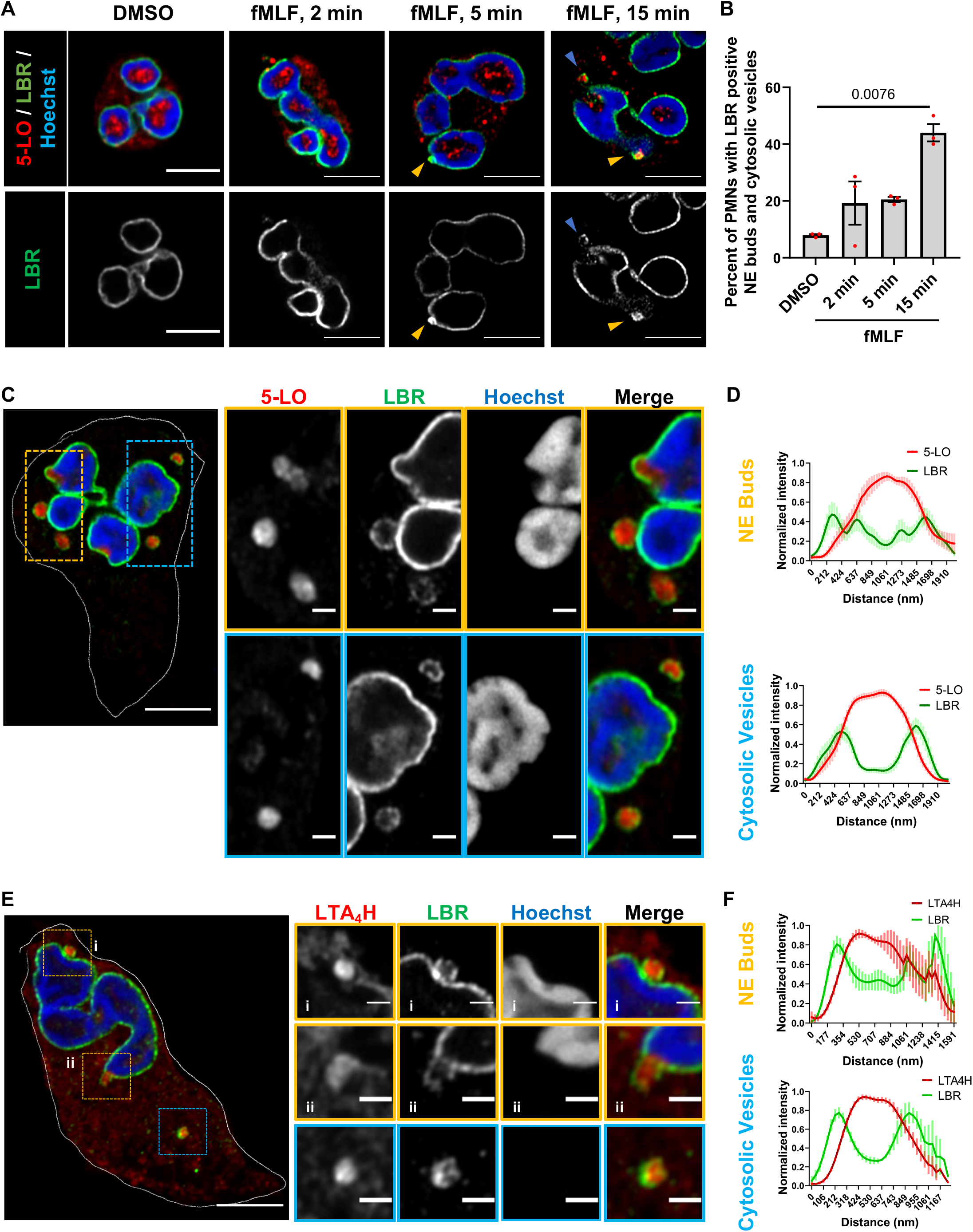
The LTB_4_-synthesizing machinery is packaged in NE-derived buds and cytosolic vesicles in activated neutrophils. **A** Representative Airyscan microscopy images showing the distribution of LBR (green) and 5-LO (red) in fixed PMNs uniformly stimulated with 20 nM fMLF (n=3). The yellow arrowheads point to nuclear buds and the blue arrowheads point to cytosolic vesicles. Scale bar is 5 µm. **B** Scatter dot plot showing the percentage of PMNs containing LBR-positive buds and cytosolic vesicles in a field of randomly selected images. Data collected from 50, 46, 74, and 66 cells in DMSO, 2min, 5 min, and 15 min fMLF treatment, respectively, pooled from three independent experiments are plotted as mean ± SEM. Each dot represents the value from one experiment. P=0.0076 as obtained from RM one-way ANOVA analysis. **C** Representative Airyscan microscopy images of PMNs chemotaxing towards 100 nM fMLF and stained for LBR (green) and 5-LO (red), obtained from six independent experiments. NE buds are shown in yellow boxes and cytosolic vesicles are shown in blue boxes. Scale bar is 5 µm. In magnified insets, the scale bar is 1 µm. **D** Histograms showing the normalized intensity of LBR and 5-LO across the diameter of LBR positive buds and cytosolic vesicles. The data points from 15 vesicles and 13 buds within 11 cells pooled from four independent experiments were plotted as mean +/- SEM, with the bold line showing the mean and bar representing the SEM. **E** Representative Airyscan microscopy images of fixed PMNs chemotaxing towards 100 nM fMLF and stained for LBR (green) and LTA_4_H (red), obtained from three independent experiments. See panel C for more details. Scale bar is 5 µm. In the inset, it is 1 µm. **F** Histograms showing the normalized intensity of LBR and LTA_4_H across the diameter of LBR positive buds and cytosolic vesicles. The data points from 5 vesicles and buds each within 5 different cells pooled from three independent experiments were plotted as mean +/- SEM, with the bold line showing the mean and bar representing the SEM.

Changes in lipid bilayer symmetry are required to induce membrane curvatures for the initiation of budding. Whereas certain transmembrane or BAR-domain-containing proteins are involved in facilitating membrane curvature, the localized presence of lipids, such as ceramide and lysophosphatidic acid, has also been reported to induce changes in membrane curvature^16^. Furthermore, lipid-ordered domains that contain ceramide and the ganglioside GM1 are required for the assembly of various signaling complexes and their endocytosis^17,18^. To assess whether membrane microdomains are involved in NE budding we performed confocal fluorescence microscopy of chemotaxing PMNs in the presence of the phase transition sensitive lipid probe di-4ANEPPDHQ^19^. We took advantage of the blue shift in fluorescence of di-4ANEPPDHQ upon binding with lipids in ordered membrane environments, to acquire general polarization (GP) images of chemotaxing PMNs, where high GP values indicate lipid ordered (L_o_) domains. Time-lapse imaging showed spatiotemporal increases in regions of high GP values in punctae close to the nucleus in PMNs chemotaxing towards fMLF (**Movie S1**) and quantification revealed the abundance of L_o_ punctae within 1µm of the nucleus compared to the rest of the cell (**Fig. 2A&B**). Sphingomyelin, the major phosphosphingolipid in mammalian cells, is hydrolyzed to ceramide and phosphocholine by the action of sphingomyelinases^20^. The enrichment of ceramide coalesces nanoscale lipid domains leading to the formation of microscopic ceramide-rich signaling platforms^21^. The nSMase1 and its substrate sphingomyelin are abundant in the nucleus^22^ and the depletion of nSMase1 from cancer cells^23^, neuronal cells^24^, or neutrophils^3^ has been reported to decrease the secretion of CD63-positive exosomes as well as FLAP/5-LO-positive exosomes from neutrophils^3^. We, therefore, assessed the role of nSMases on the generation of the nuclear lipid microdomains in chemotaxing PMNs using the nSMase inhibitor GW4869^23^ and found a strong dependence between nSMase activity and the generation of perinuclear L_o_ domains (**Fig. 2C&D**). In addition, we found similar defects in differentiated HL-60 (dHL-60) cells genetically lacking nSMase1 (**Supplementary Fig. 2A-D)**.

**Figure 2.**
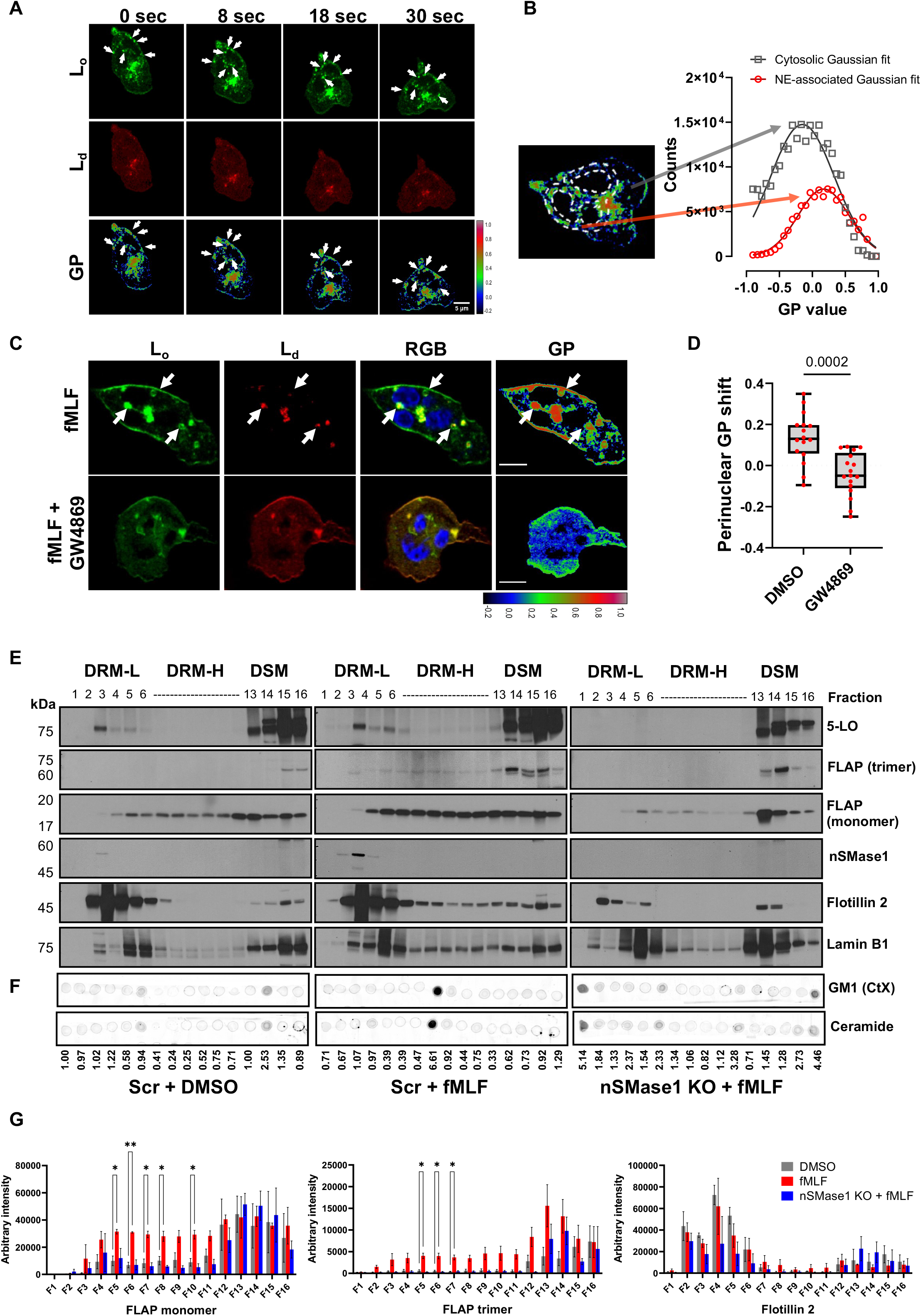
nSMase1 facilitates the recruitment of the LTB_4_ synthesizing machinery on lipid-ordered NE microdomains. **A** Representative fluorescence images of di-4-ANEPPDHQ-stained PMNs chemotaxing towards 100 nM fMLF (left) (n=5), showing the temporal distribution of the lipid-ordered (L_o_) domains (green), lipid-disordered (L_d_) domains (red), and the corresponding GP image. Pseudo-colored GP images are based on the colormap, with white/red shades depicting lipid ordered regions and blue/darker shades representing lipid disordered regions. White arrows mark the perinuclear vesicular regions of high GP values. Scale bar is 5µm. **B** Histogram depicting the gaussian distribution of the GP pixel intensity obtained from the two ROIs as depicted in the left image, with the cytosol ROI (gray arrow) 1 µm from the NE as determined by Hoechst staining (not shown) and the perinuclear ROI (red arrow) extending up to 1 µm from the NE. **C** Representative fluorescence images of di-4-ANEPPDHQ-stained PMNs chemotaxing towards 100 nM fMLF in the presence of DMSO or 3 µm GW4869, obtained from three independent experiments. The lipid-ordered (L_o_) domains (green), lipid-disordered (L_d_) domains (red), RGB images as the merge of L_d_, L_o,_ and Hoechst (blue), and GP images are presented. For color scale, refer to panel A. White arrows show the perinuclear vesicular regions of high GP values. Scale bar is 5µm. Also, see Movie S1. **D** Box-whisker plots showing the median value of the perinuclear GP intensity distribution of DMSO- and GW4869-treated PMNs chemotaxing towards fMLF. 16 data points (red dots) from the DMSO sample and 17 from the GW4869 sample pooled from three independent experiments are plotted as median with range. P=0.0002 (***) quantified using the Mann-Whitney test. **E** Representative western blots of three independent experiments, showing the distribution of various proteins in DRM-L, DRM-H, and DSM fractions isolated from the NE of Scr or nSMase1 KO dHL-60 cells stimulated with 100 nm fMLF. LaminB1 is used as a loading control. **F** Representative dot-blots of lipids isolated from the fractions shown in panel E stained for GM1 or ceramide (representative of two experiments). The ceramide intensity values of the spots relative to lane 1 of the DMSO sample are indicated below the image. **G** Bar graphs depicting the arbitrary intensity of FLAP monomer, FLAP trimer, and Flotillin 2 signal in the different fractions of either DMSO- or fMLF-treated Scr cells or fMLF-treated nSMase1 KO cells. Data from three independent experiments are plotted as mean ± SEM. P values were determined using two-way ANOVA, with DMSO as a control column, where * denotes p = 0.01-0.05 and ** depicts p = 0.001-0.01.

Lipid-ordered domains are resistant to solubilization in non-ionic detergents, by virtue of their ceramide-, GM1- and cholesterol-rich composition^25^. To assess the role of nSMase1-dependent lipid ordering in the recruitment of FLAP and 5-LO at sites of NE budding, we isolated nuclei from resting and fMLF-stimulated Scr and nSMase-1 KO dHL-60 cells and separated detergent-resistant membranes (DRMs) and detergent soluble membranes (DSMs) (**Supplementary Fig. 3A**). The purity of the nuclear preparation from resting dHL-60 cells was assessed using GAPDH (cytosol marker), calreticulin (ER marker), LBR (NE marker), Histone H3 (chromatin marker), and LTB_4_ synthesizing machinery, FLAP, and 5-LO (**Supplementary Fig. 3B**). Flotillin 2 and Lamin B1 were used as a lipid microdomain marker and as a loading control, respectively (**Fig. 2E**). We found an increase in the intensity of FLAP monomers and trimers – the functional form of FLAP^26^-along with 5-LO in both light DRM fractions (DRM-L) enriched with Flotillin 2 as well as in heavy DRM fractions (DRM-H) enriched with ceramide/GM1, upon fMLF stimulation in Scr dHL-60 cells (**Fig. 2E&F**). nSMases have been reported to bind sphingomyelin-rich nanodomains for ceramide generation and to dissociate from ceramide-rich microdomains/DRM-H^27^, accordingly, we found that nSMase1 was only enriched in DRM-L fractions (**Fig. 2E**). Remarkably, we observed a dramatic inhibition of monomeric and trimeric FLAP signals in the DRM-L and DRM-H fractions in nuclei isolated from fMLF-activated nSMase1 KO dHL-60 cells compared with Scr dHL-60 cells, which also showed a loss of ceramide/GM1 in DRM-H fractions, while Flotillin 2 levels remained grossly unchanged (**Fig. 2E-G**). Since low temperatures employed during the biochemical isolation of DRMs can artificially induce the formation of DRMs in some cases^28^, we assessed FLAP aggregation on the NE of fixed nuclei isolated from either DMSO- or fMLF-treated Scr or nSMase1 KO dHL-60 cells. As depicted in Figure 3A, we observed an increase in the regions with high FLAP signal intensity in fMLF-stimulated Scr dHL-60 cells compared to DMSO-treated Scr and fMLF-treated nSMase1 KO dHL-60 cells, demonstrating a nSMase1-dependent increase in FLAP recruitment in response to fMLF activation (also see **Movie S2**). However, the absence of nSMase1 did not alter the decrease in the nuclear sphericity observed in response to fMLF stimulation (**Fig. 3B**). We next labeled the nuclei isolated from PMNs with antibodies against ceramide and FLAP and assessed object colocalization from 3D-reconstructions of z-stacks imaged using Airyscan microscopy. We observed defects in the fMLF-induced increase in the colocalization of FLAP with ceramide clusters in the nuclei isolated from GW4869-treated PMNs (**Fig. 3C, Movie S3**). Upon quantification, we measured a significant decrease in FLAP-ceramide co-occurrence and correlation upon nSMase inhibition in fMLF activated PMNs (**Fig. 3D**). Together, these findings establish the requirement of nSMase1 for the ceramide-dependent enrichment of FLAP on the NE of activated neutrophils and dHL-60 cells.

**Figure 3.**
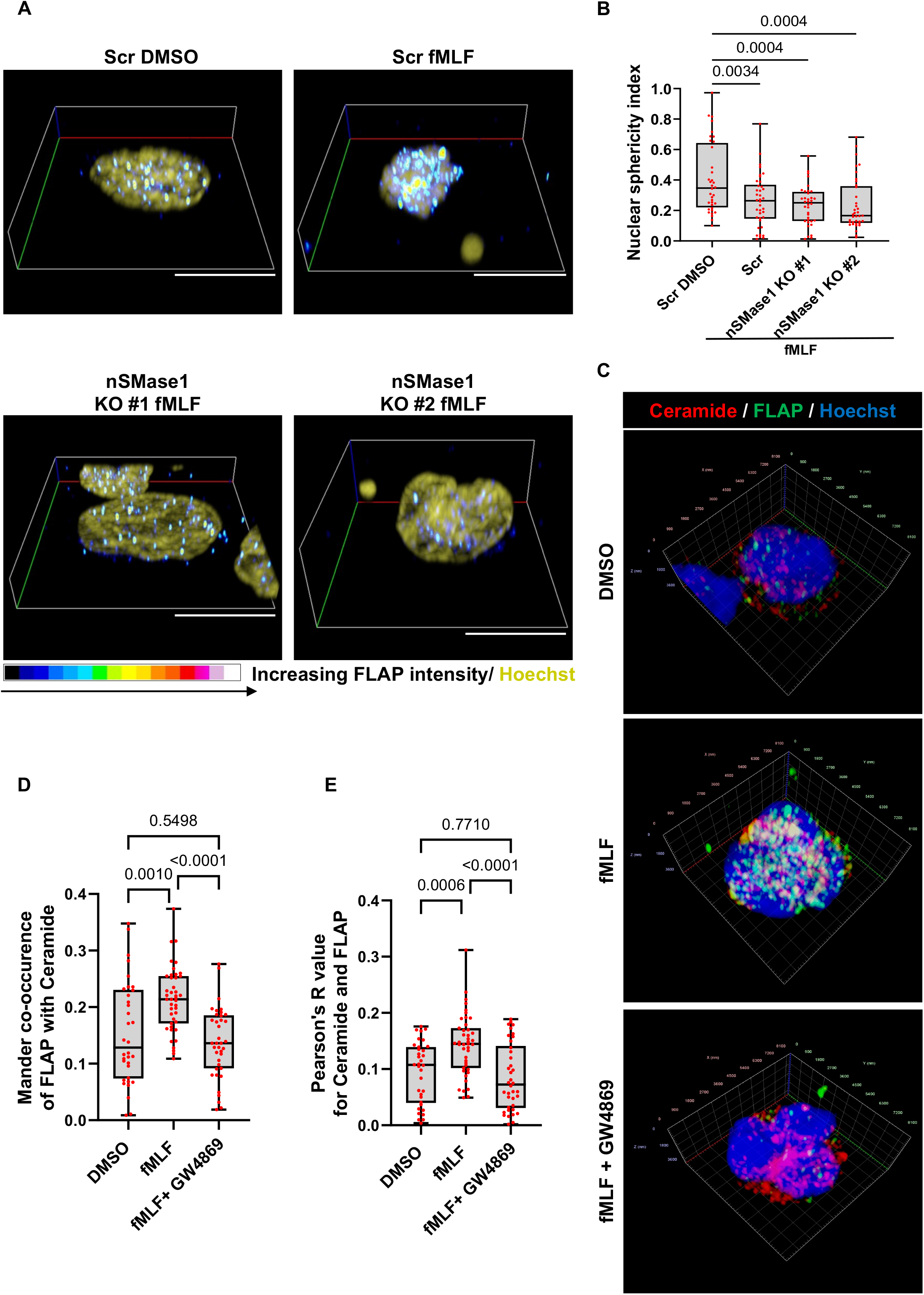
nSMase-dependent enrichment and colocalization of FLAP with ceramide-positive structures on the NE of activated neutrophils. **A** Representative 3D volumetric render of the fixed nuclei isolated from DMSO or fMLF (100 nM) stimulated Scr or nSMase1 KO dHL-60 cells stained for FLAP (spectrum) and Hoechst (olive). Red/whiter shades as shown in the colormap, denote higher FLAP intensity whereas blue/darker shades represent regions of lower FLAP abundance (n=3). Scale bar is 5 µm. Also, see Movie S2. **B** Box-whisker plots showing the sphericity of 3D-reconstructed images of isolated nuclei. Data points from at least 36 cells each pooled from three independent experiments are plotted as median with range. P values were determined using Tukey’s multiple comparisons test with ordinary one-way ANOVA. **C** Representative 3D volumetric render of the nuclei isolated from PMNs stimulated with either DMSO or fMLF (100 nM) in the presence or absence of GW4869 (3 µM), and immunostained for ceramide (red), FLAP (green), and Hoechst (blue) (n=3). Scale shown on xyz-axis. Also, see Movie S3. **D-E** Box-whisker plots showing the scatter of the Mander co-occurrence coefficient **(D)** and Pearson’s R-value **(E)** between FLAP- and ceramide-positive structures present on the NE under indicated conditions, using 3D-reconstructed multiple z-stack images. Data from three independent experiments are plotted as median with all 34 datapoints of DMSO, 43 of fMLF, and 38 of fMLF + GW4869, shown as red dots. P values determined using ordinary one-way ANOVA, are indicated on the graph.

We next sought to visualize the distribution of nSMase1 and ceramide in PMNs. We stained PMNs chemotaxing towards fMLF with ceramide and LBR and observed ceramide staining on the NE as well as on the inner periphery of LBR-positive NE buds and cytosolic vesicles (**Fig. 4A, B**). We also found a strong colocalization between nSMase1 and ceramide and between nSMase1 and 5-LO on NE buds and cytosolic vesicles (**Fig. 4C&D**). Consistent with the previously reported nuclear localization of nSMase1^22^, we observed nSMase1 localization on the NE (**Fig. 4D, green arrows**). Similarly, in dHL-60 cells expressing nSMase1-GFP chemotaxing towards fMLF, we observed a clear NE distribution along with a strong enrichment at the sites of NE buds (**Supplementary Fig. 4; Movie S4**). Since ceramide-induced changes in membrane curvature are required for the initiation of vesicle budding^29^, we investigated the effect of nSMase inhibition on NE-budding and vesiculation. We found a reduction in the percent of cells showing LBR- and 5-LO-positive NE buds and cytosolic vesicles in GW4869-treated PMNs chemotaxing toward fMLF (**Fig. 4E&F**). However, most of the few remaining LBR-positive vesicles observed in the GW4869-treated cells were NE-associated (**Fig. 4G**). These findings suggest that while other components can initiate the formation of NE buds, nSMase activity is required for the release of the NE buds and the generation of cytosolic vesicles. Indeed, the membrane bending properties of cPLA_2_^30^, which is upstream of nSMase, could contribute to the formation of NE-associated buds.

**Figure 4.**
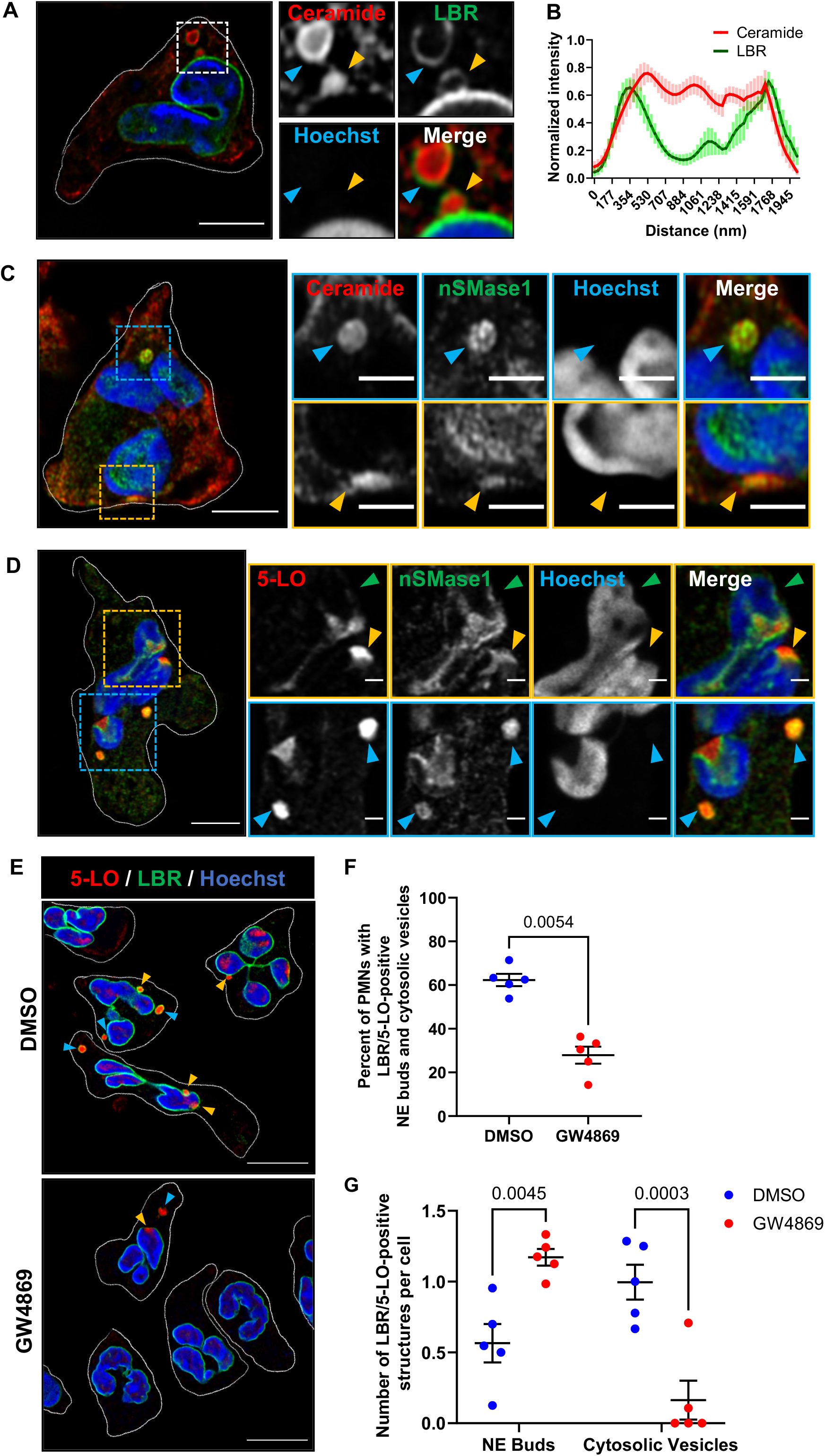
nSMase1 and ceramide are present within and are required for the generation of NE-derived 5-LO/LBR positive NE buds and cytosolic vesicles. **A** Representative Airyscan microscopy image of fixed PMNs chemotaxing towards 100 nM fMLF and stained for LBR (green) and ceramide (red) (n=4). The yellow arrowheads point to nuclear buds and the blue arrowheads point to cytosolic vesicles. Scale bar is 5 µm. In the inset, it is 1 µm. **B** Histogram showing the normalized intensity of LBR and ceramide across the maximum width of the LBR-positive vesicles. Data are represented as mean ± SEM of 10 NE buds or cytosolic vesicles from three independent experiments. **C** Representative Airyscan microscopy images of fixed PMNs chemotaxing towards 100 nM fMLF and stained for nSMase1 (green) and ceramide (red) (n=3). The yellow arrowheads point to nuclear buds and the blue arrowheads point to cytosolic vesicles. Scale bar is 5 µm, in the inset it is 2 µm. **D** Representative Airyscan microscopy images of fixed PMNs chemotaxing towards 100 nM fMLF stained for 5-LO (red) and nSMase1 (green) (n=3). The yellow arrowheads point to nuclear buds and the blue arrowheads point to cytosolic vesicles. Green arrowheads point to the 5-LO and nSMase1 on the NE. Scale bar is 5 µm. In the inset it is1 µm. **E** Field Airyscan microscopy image of fixed PMNs chemotaxing towards 100 nM fMLF in the presence or the absence of 3 µM GW4869 stained for 5-LO (red) and LBR (green). Scale bar is 10 µm. The yellow arrowheads point to nuclear buds and the blue arrowheads point to cytosolic vesicles. **F** Scatter plots showing the percentage of PMNs in DMSO- or GW4869-treated PMNs containing LBR/5-LO-positive NE buds and cytosolic vesicles among the total cells quantified within randomly selected imaging fields. Data points from five independent experiments (red dots) are plotted as mean ± SEM. A total of 158 cells in the DMSO sample and 108 cells in GW4869 samples, were analyzed. P values determined using a two-tailed paired t-test are reported. **G** Scatter plots showing the changes in the number of LBR/5-LO-positive buds and cytosolic vesicles per cell in PMNs treated with GW4869 compared to DMSO control. The data points from five independent experiments (red dots) are plotted as mean ± SEM, and P values determined using two-way RM ANOVA are reported.

To further characterize the nature of the 5-LO/FLAP/LBR-positive NE buds and cytosolic vesicles, we used expansion microscopy – a method developed to enable confocal microscopy to visualize sub-diffraction limited details by isotropic enlargement of the samples^31,32^. PMNs chemotaxing under agarose were fixed, stained, crosslinked, and expanded as described in the methods section, giving rise to an expansion of the sample by ∼4x. We observed the clear presence of 5-LO-positive punctae within the LBR-positive cytosolic vesicles **(Fig. 5A)** and NE-associated buds **(Fig. 5B)** in chemotaxing cells. We also noted the presence of ceramide mainly at the periphery of the LBR-positive structures, although small ceramide punctae could also be seen inside the structures (**Fig. 5A-B**), as observed using traditional IF imaging (**Fig. 4A-B)**. The median diameter of the LBR-positive NE buds and cytosolic vesicles was 972.5 nm (**Fig. 5C**), which closely matched the sizes measured using conventional confocal imaging (1000 ± 200 nm) (**Supplementary Fig. 1B**), and the median diameter of the 5-LO positive punctae present within the LBR NE buds or cytosolic vesicles was 200 nm (**Fig. 5D**). We also observed a positive correlation between the size of the LBR-positive vesicles and the number of 5-LO-positive punctae within the LBR-positive vesicles (**Fig. 5C**). Notably, we did not observe the presence of the canonical exosome marker CD63 within the LBR-positive structures, although CD63-positive structures did contain ceramide punctae, indicative of ceramide enriched conventional ILVs^33^ (**Fig. 5E&F, Supplementary Fig. 5A&B**). The CD63-positive vesicles had a smaller diameter that ranged between 350-420 nm (**Supplementary Fig. 5C**), which matches the reported size range of CD63-positive MVBs^34^. Using traditional IF, we also found that the number and size of the CD-63-positive vesicles did not change in response to fMLF stimulation (**Supplementary Fig. 5D&E**). We previously observed the colocalization of mCherry-5-LO with CD63-GFP in chemotaxing dHL-60 cells^3^. We envision that this was a consequence of 5-LO overexpression and/or to the potential multimerization from the mCherry fusion, targeting mCherry-5-LO to the CD63 positive vesicles. Together, these findings show that the LBR-positive MVBs contain 5-LO-positive ILVs and are distinct from the canonical CD63-positive MVBs in both their composition and their size.

**Figure 5.**
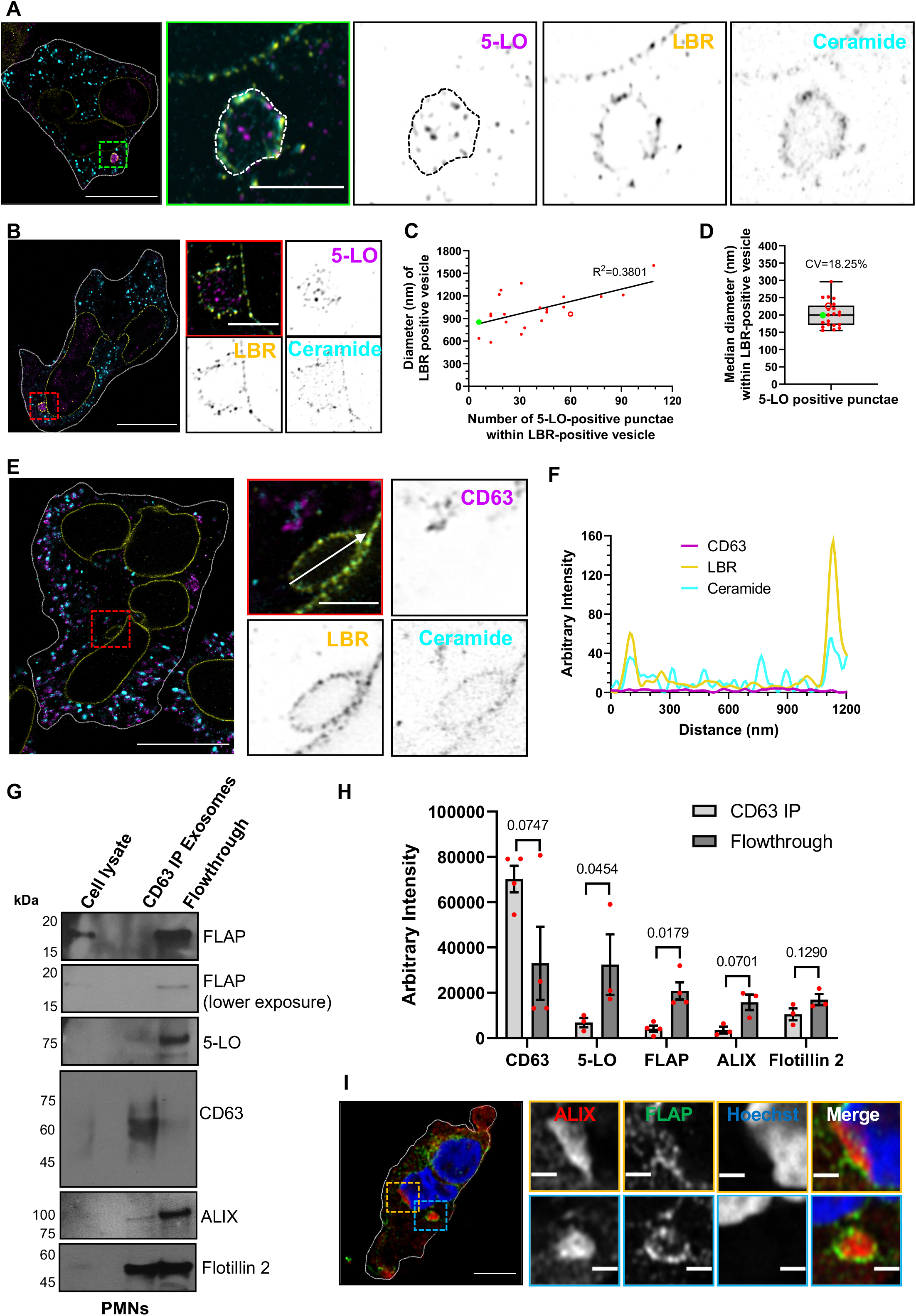
5-LO-positive and CD63-negative punctae are present within LBR-positive vesicles. **A-B** Four-fold expansion images of fixed human PMNs chemotaxing towards 100 nM fMLF, acquired using Airyscan microscopy, stained for 5-LO (magenta), LBR (yellow), and ceramide (cyan) within cytosolic vesicles **(A)** and NE buds **(B)**, are represented as inverted grayscale zoomed images (n=3). Scale bar is 5 µm. In the inset, the scale bar is 800 nm. **C** Scatter plot showing the non-linear regression of the diameter of LBR positive vesicles versus the number of 5-LO positive punctae per LBR-positive vesicle. **D** Box-whisker plot showing the distribution of the median diameter of 5-LO-positive punctae per LBR-positive vesicle. Data are plotted as median with range of 22 LBR-positive vesicles (red dots) within 10 cells pooled from two independent experiments. In panels C and D, the blue dot and open red dot represent data points from panels A and B, respectively. **E** Four-fold expansion microscopy image of fixed human PMNs chemotaxing towards 100 nM fMLF, acquired using Airyscan microscopy, stained with CD63 (magenta), LBR (yellow), and ceramide (cyan) within an NE-derived vesicle, represented as inverted grayscale zoomed images (n=3). Scale bar is 800 nm. The scale bar in the uncropped image is 5 µm. **F** Histogram showing the arbitrary intensity of LBR, ceramide, and CD63 across the white arrow along the LBR positive bud shown in zoomed and merged image in panel E. **G** Representative western blot of exosomes obtained from fractions 4-9 of the density gradient centrifugation from PMNs stimulated with 100 nM fMLF and immunoprecipitated using an antibody against CD63 (n=3-4). CD63-positive and CD63-negative (unbound flow-through) populations were immunoblotted for FLAP, 5-LO, CD63, ALIX, and Flotillin 2. **H** Bar graph showing the quantifications of the band intensity of CD63, 5-LO, FLAP, ALIX, and Flotillin 2 in CD63-IP and flowthrough fractions. Four data points for FLAP and CD63, and 3 for 5-LO, ALIX, and Flotillin 2 are plotted as mean ± SEM where each red dot represents the value from one experiment. P values determined using a ratio paired t-test are reported. **I** Representative Airyscan microscopy images of fixed PMNs chemotaxing towards 100 nM fMLF and stained for FLAP (green) and ALIX (red) (n=3). Scale bar is 5 µm. In the inset, the scale bar is 1 µm. NE buds are shown in yellow boxes and cytosolic vesicles are shown in blue boxes.

As we previously reported using shRNA mediated nSMase1/SMPD2 knockdown HL-60 cells^3^, we found that fMLF-stimulated nSMase1 KO dHL-60 cells release fewer exosomes, compared to Scr dHL-60 cells. Using nanoparticle tracking analysis (NTA), we found that Scr cells released two major populations of exosomes in response to fMLF stimulation (**Supplementary Fig. 6A&B**). Interestingly, exosomes isolated from nSMase1 KO dHL-60 cells showed a significant decrease in the number of the larger, >180 nm, particles, which correlated with the size of the 5-LO punctae within the LBR-positive MVBs measured using expansion microscopy (**Fig. 5D**). We also observed a decrease in the levels of 5-LO and FLAP (**Supplementary Fig. 6C&D**) and of LTB_4_ content (**Supplementary Fig. 6E**) in exosomes isolated from the nSMase1 KO cells. Although exosomal Flotillin 2 levels did not change significantly in the nSMase1 KO cells, CD63 levels decreased (**Supplementary Fig. 6C&D**), as we previously published^3^. A closer analysis of the fractions of the supernatants from fMLF-stimulated PMNs separated by density gradient ultracentrifugation revealed that the levels of the canonical exosome markers CD63, Flotillin 2, and TGS101 peaked in higher density fractions compared with FLAP and the ESCRT-associated protein ALIX^9^, which appeared more uniformly distributed throughout fractions 4-9 (**Supplementary Fig. 7A&B**). To further assess this, we used anti-CD63 antibody immunoprecipitation of the exosomes pooled from fractions 4-9 of fMLF activated PMNs and compared the composition of CD63-positive and flowthrough exosome populations. We found that while Flotillin 2 was present in both CD63-positive and flowthrough exosome preparations, CD63-positive exosomes showed decreased FLAP and 5-LO signal (**Fig. 5G&H**). Moreover, although ALIX was present in CD63-positive exosome pulldowns, it was enriched in the FLAP/5-LO-positive exosome elution (**Fig. 5G&H**). These results are supported by the exclusion of CD63 signal from LBR-positive NE buds and cytosolic vesicles (**Fig. 5E&F, Supplementary Fig. 5D**) and the substantial recruitment of ALIX to LBR- or FLAP-positive NE buds and cytosolic vesicles in PMNs chemotaxing towards fMLF (**Fig. 5I, Supplementary Fig. 8**). From these findings we conclude that the NE-derived and nSMase1-dependent 5-LO- and FLAP-positive exosomes are synthesized in a CD-63 independent pathway.

Neutrophil infiltration and exit into and from sites of infection/injury are crucial for potent inflammatory response and tissue homeostasis. Neutrophils are endowed with a highly malleable multilobed nucleus enriched with LBR and expressing high levels of Lamin B1 and relatively low levels of Lamin A/C^15^. This unique nuclear architecture has been implicated in neutrophil extravasation and squeezing through tight spaces during migration^35^. The LTB_4_ signaling pathway plays a key role during neutrophil chemotaxis *in vitro*^3^ and *in vivo*^10^ as well as during extravasation^36,37^. We now show that the biogenesis of exosomes containing the LTB_4_-synthesizing machinery originates at sites of NE budding during neutrophil chemotaxis. Our study identifies nSMase1 as being critical for the generation of NE-derived buds and cytosolic vesicles and the subsequent release of FLAP/5-LO containing exosomes. More importantly, we report that nSMase1 localization on the NE is required for ceramide production, which leads to the generation of lipid-ordered nuclear membrane microdomains and the recruitment of FLAP/5-LO at the sites of NE buds. The induction of membrane curvature required for vesicle budding is generally mediated by (i) lipid composition, (ii) protein motif insertion, and/or (iii) clustering of membrane proteins of defined shapes^16^. Ceramide and its glucoside derivative, GM1 are known to facilitate membrane budding by inducing negative and positive membrane curvature, respectively^38,39^. The enrichment of these lipids at sites of nuclear budding underscores their role in the initiation of FLAP/5-LO-containing NE buds. Moreover, the induction of positive membrane curvature mediated by the insertion of the C2-domain of cPLA_2_ and lysoPC, the byproduct of cPLA_2_ activity^30^, may facilitate the budding of the vesicle from the NE. Indeed, cPLA_2_ was recently proposed to act as a molecular sensor of nuclear membrane tension in migrating cancer cells and zebrafish^40,41^. Notably, the released AA through cPLA_2_ activation is a key factor in LTB_4_ biogenesis, where it was shown to not only induce FLAP trimerization^42^ and act as the substrate for 5-LO but also as an activator of nSMases and ceramide production^43^. We, therefore, propose that the clustering of FLAP, along with the incorporation of ceramide, induces membrane curvatures that are required for NE-derived bud formation. However, as we observe the presence of the INM protein LBR on 5-LO- and FLAP-positive cytosolic vesicles, the transformations needed to generate an MVB from the two NE membranes remain unknown. Interestingly, the emergence of MVBs from the NE has previously been reported in other cell types^44,45^, where the MVBs appear to be enclosed in two membranes. Since ALIX binds and sorts the membrane-associated tetraspanin cargo to ILVs/exosomes^46^, the presence of ALIX within FLAP-positive NE buds and cytosolic vesicles in chemotaxing PMNs also suggests a FLAP-mediated recruitment of ALIX at sites of NE buds and ILV invagination. Furthermore, ALIX recruits the ESCRT III complex protein CHMP4B, which is known to induce membrane curvature during vesicle fission and ILV formation^47^. Additional studies are needed to determine the mechanism underlying the fission of NE membrane and ILV generation. Finally, as we found that the absence of nSMase1 significantly downregulates the secretion of a larger (>180 nm) 5-LO-positive exosome population that originates within LBR-positive and CD63-negative MVBs, we propose that activated neutrophils release at least two structurally, biochemically, and functionally distinct exosomes populations: namely NE-derived and conventional exosomes. Indeed, exosome heterogeneity based on size/density^48^, cargo type^49^, and the mechanism of ILV generation^50^, has recently been reported. We envision that the unique characteristics for the neutrophil nuclei provide a specialized environment for the release of nuclear material during chemotaxis.

## ACKNOWLEDGEMENTS

We thank the Platelet Pharmacology and Physiology Core at the University of Michigan for providing human blood from healthy volunteers. We are grateful to Drs. Dawen Cai and Ye Li (University of Michigan) for their assistance in the expansion microscopy experiments. We also acknowledge Drs. Ritankar Majumdar and Cosmo Saunders as well as Jeannie Yoojin Park for their intellectual contributions. Finally, many thanks to members of the Parent laboratory, and Drs. Phyllis Hanson (University of Michigan) and Pierre Coulombe (University of Michigan) for their valuable suggestions and help during the progress of this work. This work was supported by funding from the University of Michigan School of Medicine (CAP), by the Lucchesi Predoctoral Fellowship Award (SC), by an American Heart Association Predoctoral Award (FJ), and by R01 AI152517 (CAP).

## ONLINE METHODS

### Isolation of human peripheral blood neutrophils

Heparinized whole blood from anonymous healthy human donors that had not taken aspirin for 7 days and NSAIDS for 48 hours was obtained by venipuncture from the Platelet Pharmacology and Physiology Core at the University of Michigan. Neutrophils were isolated using the protocol described earlier^51^. Briefly, whole blood was incubated with an equal volume of 3% dextran (sigma D1037) in 0.9% NaCl for 30 min at 37°C to facilitate RBC sedimentation. Three volumes of plasma containing platelets monocytes, lymphocytes, and neutrophils were overlayed onto a volume of Histopaque-1077 (Sigma 10771) and centrifuged at 400Xg for 20 min at room temperature to separate peripheral blood mononuclear cells from neutrophils. Residual erythrocytes in the neutrophils pellet were removed using ACK lysing buffer (Thermo Fisher A1049201). The protocol yields >99% live neutrophils with >95% purity.

### Cell lines and plasmid constructs

The human myeloid leukemia-derived pro-myelocytic cell line HL-60 was obtained from ATCC and maintained in RPMI-1640 media containing 10% HI-FBS, 20 mM HEPES pH 7.4, and penicillin-streptomycin antibiotic cocktail. To generate neutrophil-like cells HL-60 were differentiated in culture media containing 1.3% DMSO for 6 days with a change to fresh media every other day as described by Saunders *et al* ^52^. HEK293T cells cultured in DMEM containing 10% FBS were used to generate lentiviral particles for the generation of stable HL-60 cell lines. pVSVG, pCMV dR8.91, and pLentiCRISPR V2 vector expressing SCR/nSMase1 sgRNA or pCDH MSCV MCS EF1 neomycin vector expressing nSMase1 fused with eGFP at the C-terminal were transfected to HEK293T at the ratio of 1:2:4 using Lipofectamine 3000 transfection reagent. The lentiviral particles collected after 48- and 72-hours post-transfection were pooled, concentrated using PEG-it (Systems Biosciences LV810A-1), and added to the HL-60 cells with 8ug/ml polybrene. The clones expressing the construct were selected in 2 µg/ml puromycin and verified by western blotting and genetic sequencing. The SMPD2 sgRNA #1; GCCGACCGCATGAGGCGCCT, and sgRNA#; GAACCAGGAGAGCTTCGACC were cloned in pLentiCRISPR V2 plasmid, which was a kind gift from the Zhang lab. Full-length SMPD2 (NM_003080) was amplified from the cDNA pool generated using oligo dT primer based SuperScript™ IV First-Strand Synthesis System (Thermo Fischer 18091050) kit, using RNA extracted from dHL60 cells. Gene-specific primers: 5’ GGAATTCGCCACCATGAAGCCCAACTTCTC 3’ and 5’ CCGCTCGAGTTGTTCTTTAGTTCTGTCC 3’, were used to amplify SMPD2 with 5’ EcoRI and 3’ XhoI RE sites. The fragment was cloned in pCDH MSCV MCV EF1 Neomycin vector upstream to 5’ XhoI-EGFP using T4 DNA ligase.

### Isolation of intact nuclei for microscopy and the purification of nuclear membrane microdomains

The protocol is based on and modified from the DRM isolation protocols by Persaud-Sawin *et al*.^53^ and Cascianelli *et al*.^54^, as is shown in Supplementary Figure 3. dHL-60 or PMNs were resuspended in 1X mHBSS (150 mM NaCl, 4 mM KCl, 1.2 mM MgCl_2_, 10 mg/ml Glucose, 20 mM HEPES pH 7.4) at a density of 10e6/ml. The cells were treated with either DMSO or 100 nM fMLF (Sigma Aldrich F3506) in the presence or absence of 3 µM GW4869 (Sigma Aldrich D1692) for 5 min at 37°C with 10 RPM rotation. The cells were further incubated with an equal volume of 20 mM dimethyl pimelimidate (DMP) crosslinker (TCI chemicals D4476) in 1X mHBSS for 15 min at 37°C, washed in 20 mM Tris-Cl (pH 8.0) at 500Xg for 5 min at 4°C, and partially lysed twice with 15X trituration in ice-cold hypotonic lysis buffer (10 mM HEPES pH 7.5, 4 mM MgCl_2_, 25 mM NaCl, 1 mM DTT, and 0.1% NP-40) at a density of 50e6/ml. After centrifugation at 16,000Xg at 4°C for 10 sec, supernatants were saved as vesicle containing cytosol fractions. To remove ER fragments (microsomes), Golgi, and mitochondria, the nuclear pellet was washed with ice-cold Barnes solution (85 mM KCl, 85 mM NaCl, 2.5 mM MgCl_2_, 5 mM trichloroacetic acid pH 7.4)^55^. For immunostaining, the purified nuclei were resuspended in ice-cold isotonic resuspension buffer (10 mM HEPES pH 7.4, 4 mM MgCl_2_, 150 mM NaCl, 1 mM DTT, 250 mM Sucrose) at a density of 1e6/ml for microscopy, added to the poly-L-lysine coated glass coverslip, spun at 500Xg for 5 min at 4°C and fixed with 4% paraformaldehyde in isotonic resuspension buffer for 15 min at room temperature. For the isolation of lipid microdomains, nuclei were further lysed in 650 µl ice-cold TNE buffer (50 mM tris-Cl pH 7.4, 150 mM NaCl, 5 mM EGTA) containing 4 mM MgCl_2_ and 1% Triton X-100. The suspension was passed through a 23 G needle 30X and incubated on ice for 30 min. The NE supernatant was collected at 110Xg for 10 min at 4°C and adjusted to 40% iodixanol concentration using 60% Optiprep solution (Sigma D1556) and overlayed with 7 ml of 30% iodixanol, 2 ml of 20% iodixanol, and 1 ml of 5% iodixanol solution in TNE buffer, in a 13 ml ultracentrifuge tube. Sixteen fractions of 750 µl each from the top (F1) were collected after centrifugation at 150,000Xg for 16 hrs at 4°C. One-fifth of the volume of the individual fraction was used to extract lipids by the methanol-chloroform method as described by Moltu *et al* ^56^. The rest of the fraction volume was used to concentrate protein by the TCA-acetone method, and proteins were resuspended in 75 µl of 1X Laemmli sample buffer and boiled at 95°C for 10 min before loading on 4-12% Bis-Tris gel for electrophoresis. The electrophoresed proteins were transferred to 0.2-micron PVDF membrane, blocked using 1X Fish gelatin (Fischer Scientific NC0382999) in TBS containing 0.1% Tween-20 and probed for specific proteins using antibody against FLAP (1 µg/ml, Abcam 85227), 5-LO (1:1000, Abcam 169755), Flotillin 2 (1:1000, CST 3436), nSMase1 (1:500, CST 3867), and Lamin B1 (1:1000, Abcam 133741). The lipids were spotted on the nitrocellulose membrane, blocked with 1X fish gelatin in DPBS and dot blots were probed with 1 µg/ml CF^®^568-conjugated cholera toxin (Biotium 00071) in DPBS for GM1 and with mouse anti-ceramide antibody (1:500, Sigma C8104-50TST) followed by the Texas red-conjugated anti-mouse IgM secondary antibody to detect ceramide.

### Underagarose chemotaxis assay, live imaging, and immunofluorescence microscopy

The chemotaxis assay was performed as described by Saunders *et al* ^52^. Cell culture dishes were coated with 1% BSA in DPBS at 37°C for 1 hr. 0.5% agarose in DPBS: HBSS (1:1) was poured and allowed to solidify for 45 min. Three 1 mm diameter wells were carved 2 mm from each other. fMLF (100 nM) in HBSS was placed in the middle well creating a gradient of 50 pM/µm as described by Afonso and colleagues^5^. A total of 50,000 cells in 5 μl mHBSS were plated in the outer wells and incubated at 37°C. The chemotaxing cells were imaged under a 63X oil objective in a temperature-controlled chamber set at 37°C. Images of nSMase1-GFP expressing HL-60 cells were acquired at 10-sec intervals, using Airy disk imaging array “Airyscan” to rapidly achieve resolution beyond the diffraction limit (140 nm at 488 nm). The acquired images were reconstructed using the Airyscan processing tool using the Zen imaging software and were converted to a movie using Fiji. For immunostaining, the PMNs were allowed to chemotax for 1 hr and fixed with 4% PFA in HBSS for 20 min at 37°C. The agarose was removed and PMNs were blocked for 1 hour at room temperature followed by staining in blocking buffer (0.2% saponin, 2% goat serum in 1X mHBSS) at 4°C overnight with antibodies against FLAP (1µg/ml, Abcam 85227), 5-LO (1:500, BD biosciences 610694), nSMase1 (1:100, Abcam 131330), and Lamin B1 (1:100, Proteintech 66095-1), LBR (1:400, Abcam 32535), LTA_4_H (1:100, Santacruz biotechnology sc23070), Ceramide (1:50, Sigma C8104-50TST), CD63 (1:800, BD biosciences 556019), and ALIX (1:200, Abcam 117600). The PMNs were washed with HBSS 3X for 5 min each and incubated with Alexa fluor-conjugated secondary antibody along with 1 µg/ml of Hoechst 33342, for 1 hr at room temperature. The washed PMNs were mounted with Immumount™ (Fischer Scientific FIS9990402), z-stacks were acquired at an interval of 160 nm, using 63x objective in Zeiss LSM 880 confocal microscope fitted with Airyscan, and the acquired images were reconstructed using Airyscan processing in Zen 3.4 Blue edition software. The immunostaining of isolated fixed nuclei was performed in non-permeabilizing conditions (2% goat serum in DPBS).

### di-4ANEPPDHQ fluorescence imaging

The image acquisition and anisotropy quantification were performed as described earlier by Owen *et al*^19^. Briefly, cells were incubated with HBSS containing 1 µM di-4ANEPPDHQ (ThermoFisher D36802) and Hoechst for 30 min at 37°C, washed, and added to the under agarose well as described above. Cells chemotaxing towards fMLF were imaged using a 63X oil immersion lens at 37°C. di-4NEPPDHQ fluorescence was excited using 488 nm argon laser, and the emission was collected at 500–580 nm (L_o_) and 620–750 (L_d_) nm range, respectively. Hoechst was excited using a 405 nm laser and the signal was captured using a simultaneous line scanning approach on a Zeiss 880 Airyscan microscope. Images were captured at 10-sec intervals and reconstructed by Airyscan processing. Generalized polarization (GP) images were created based on the methodology outlined by Owen *et al*^19^. To calculate the perinuclear GP shift, the brighter lipid ordered (L_o_, green) channel was used to create a mask of the whole cell and the perinuclear mask with a width of 15 pixels was created using Hoechst image. The cell cytosol area was obtained by subtracting the perinuclear mask from the whole-cell mask. The GP value of every pixel in both the perinuclear area and the cytosol area was calculated using the formula below:

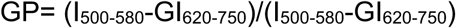

Where I_500-580_ and I_620-750_ are pixel intensities acquired in the L_o_ and L_d_ emission channels and G is the calibration factor manually adjusted to center the GP histogram of cytosol area around zero. GP distributions were obtained from the binned GP values and fitted into non-linear Gaussian using GraphPad Prism. Finally, after fitting the GP histograms of the cytosol area and perinuclear area, the GP shift = the median value of GP perinuclear histogram - the median value of GP cytosol histogram.

### Exosome Isolation, immunoprecipitation, and LTB_4_ ELISA

Exosome isolation was performed following the guidelines described by Thery *et al*^57^. PMNs or Scr and nSMase1 KO HL-60 cells were stimulated with 100 nM fMLF in RPMI-1640 containing 10 U/ml DNaseI (Sigma Aldrich DN25) for 30 min at 37ºC and the supernatants were collected at 500Xg at 4°C for 5 min. The microvesicles and apoptotic bodies were removed at 4000Xg for 20 min followed by filtration through a 0.2 µm polyethersulfone membrane filter. The extracellular vesicles (EVs) present in the filtered supernatant were concentrated with 8% PEG-6000 (Bio Basic PB0432) in 20 mM HEPES and 500 mM NaCl, at 4°C for 36 hrs, followed by centrifugation at 4000Xg at 4°C for 1 hr. The concentrated EVs, resuspended in 1 ml 250 mM sucrose and 20 mM Tris-Cl pH 7.4, were overlayed on the top of optiprep gradients and centrifuged at 100,000Xg for 16 hrs at 4°C. The optiprep gradients were prepared in 250 mM sucrose and 20 mM Tris-Cl, starting from the bottom as 3 ml of 40% optiprep, 3 ml of 20% optiprep, 3 ml of 10% optiprep, and 2 ml of 5% optiprep. The fractionated exosomes were collected as 12 fractions of 1 ml each from the top (lower to higher density).

For exosome immunoprecipitation, fractions 4-9 (Iodixanol density 1.083-1.142 g/ml) as described in Majumdar *et al*.^3^ were pooled, diluted in DPBS, centrifuged at 100,000Xg for 1 hr, resuspended in DPBS containing 1% 0.2 µm filtered BSA, and incubated with anti-CD63 antibody-conjugated magnetic beads (Thermo Scientific 10606D) overnight at 4°C with 10 RPM rotation. The CD63-specific bead-bound exosomes were separated using DynaMag™ magnets (Thermo Scientific 12321D) and washed once with DPBS containing 1% BSA. The supernatant containing unbound exosomes was collected and centrifuged at 100,000Xg for 1 hr. The unbound exosome pellet and the CD63 positive bead-bound exosomes were lysed in RIPA buffer for 15 min on ice, boiled in an equal volume of 2X reducing Laemmli buffer at 95°C for 5 min, loaded in 4-12% bis-tris gel, electrophoresed, and transferred to PVDF membrane for western blotting with anti-ALIX antibody (1:1000, Abcam 117600), anti-CD63 antibody (1:500, BD biosciences 556019), anti-Flotillin 2 antibody (1:1000, cell signaling technology 3436), anti-5-LO antibody (1:1000, Abcam 169755) and anti-FLAP antibody (1µg/ml, Abcam 85227), and detected using protein A-HRP (1:5000, Invitrogen 101023).

The LTB_4_ ELISA kit (Cayman Chemicals 520111) was used to assess the LTB_4_ levels within the exosomes. The isolated exosomes were homogenized in 100 µl ELISA buffer using a 3 mm diameter sonicator probe at an amplitude of 20% with 2 sec ON/OFF cycles for a total of 10 cycles on ice. To detect LTB_4_ concentrations within the linear range, 50 ul of concentrated homogenate was diluted 4X in ELISA buffer and LTB_4_ levels were quantified according to the manufacturer’s instructions. The values obtained were plotted using GraphPad prism.

### Nano-tracking analysis

The data for NTA was captured using a Malvern Nanosight NS300 equipped with a 488 nm laser and a high sensitivity sCMOS camera and analyzed using the NTA 3.3 Dev Build 3.3.301 software. The exosomes purified using optiprep-density gradient centrifugation were resuspended in DPBS, vortexed, and diluted to 1:1000 in 0.22 µm filtered particle-free water to obtain a recommended concentration range of 1-10 × 10^8^ particles/ml for reliable measurement. Using a syringe pump speed of 100/AU to inject exosome suspension in the flow channel, videos of the particle’s inflow were captured in script control mode, as 5 videos of 60 s each with 1 s delay and viscosity of water at 25°C. A total of 1,500 frames/sample at a capture rate of 25 frames/sec at constant camera level for each experimental set were captured.

### Expansion microscopy

To maintain the structural integrity of PMNs, samples were processed with a few modifications from the protocols described earlier^31,58^. PMNs migrating under 3 ml of agarose over a 22×22 mm glass coverslip (#1.5, BSA coated) in a 35-mm dish were fixed with 1 ml of 4% PFA and 0.05% glutaraldehyde in PHEM buffer (60 mM PIPES, 25 mM HEPES, 10 mM EGTA and 2 mM MgCl_2_, pH 6.9) at 37ºC for 20 min. After careful removal of the agarose, the fixed cells were immunostained in saponin-containing buffer, as described above. Post staining, the cells on the coverslips were washed thrice with 1X PBS and crosslinked with 3 mM Acryloyl-X, SE (Sigma Aldrich A20770) in 1X PBS, overnight at room temperature. To remove residual cross-linker, cells were washed thrice with 1X PBS for 15 min each at room temperature. The coverslips were incubated with an 80 µl drop of gelation solution comprising 8% sodium acrylate (Sigma Aldrich 408220), 10% Acrylamide (Sigma Aldrich 4058), 0.1% bisacrylamide (Sigma Aldrich M1533), 2M NaCl, and 1X PBS containing 1% heat-induced initiator VA-044 (Fischer scientific A3012), at 4ºC for 10 min. The coverslip was assembled in the gelation chamber as illustrated by Truckenbrodt *et al*.^58^, 200 µl of gelation solution was added, and the gelation chamber was incubated in a humidified chamber at 37ºC for a minimum of 2 hrs to allow polymerization. The gelation chamber was disassembled and the gel on the coverslip was washed quickly with 1X PBS to remove the unpolymerized gel. The gel was treated with 2 ml of the digestion buffer consisting of 50 mM Tris-Cl, 800 mM Guanidine hydrochloride, 1 mM EDTA, and 0.5% Triton-X 100 adjusted to pH 8.0, with 8 U/ml of proteinase K (NEB P8107S) added just before use, in a 35 mm dish, at 37ºC for 1 hr. The gel containing the homogenized sample was washed thrice with 1X PBS (2 ml each) for 15 min each at room temperature. The gel was slid carefully into a 100 mm dish to allow for gel expansion, by sequentially changing the buffer to 10 ml of 1X>0.5X>0.02X>0.01X PBS for 20 min each at room temperature, followed by incubation in ddH_2_O overnight at 4ºC. Gel size both before and after the expansion was measured using a ruler and was found to be approximately 4X lengthwise. Gel pieces containing the cells were excised to fit into a 12 mm glass-bottom 35 mm dish coated with poly-L-lysine (Sigma Aldrich P8920). The gel was gently pressed using a soft painting brush to ensure its adherence to the coverslip and avoid gel drift during imaging. 100µl ddH2O was slowly added to the empty spaces of the chamber to cover the gel and avoid the shrinking of the polymer. The sample was imaged using a Zeiss 880 confocal microscope fitted with an Airyscan detector, under a 63X (1.4NA) oil objective. Since Airyscanning provides 120 nm lateral and 350 nm axial resolution^59^, using 4X expansion of the sample has increased the resolution between 30-40 nm laterally and approximately 100 nm axially. The z-stacks acquired at an interval of 160 nm were ∼50 nm apart, after expansion adjustment. All the scales shown in the images were after 4X adjustment, as verified by measuring the CD63 positive MVB size (∼400 nm post adjustment). All the zoomed images representing buds/vesicles were deconvoluted using fast iterative algorithm and were used for presentation and data quantification.

### Image quantification and data representation

Nuclei sphericity as presented in figure 3B, was quantified from z-stacks of Hoechst-stained nuclei. Briefly, the entire stack is converted to 8-bit and intensity thresholded (Otsu) with dark background for individual images. Volume, surface area, and sphericity were calculated using analyze>analyze 3D options of MorphLibJ plugin. To quantify the colocalization between ceramide and FLAP in the nuclei, 3D images were thresholded using maximum entropy parameters, and Pearson’s R-value and Mander colocalization coefficients (figure 3D) were determined using the JaCoP plugin in Fiji image analysis software. Intensity profiles of RGB channels (Figures 1D, 1F, 4B, 5F, and supplementary figure 5B) across the diameter of the vesicle of interest were determined using the RGB profiler plugin (https://imagej.net/plugins/rgb-profiler) from the Fiji image analysis tool. The resulting data was exported to GraphPad prism and plotted as histograms with mean ± SEM. The size of LBR-positive vesicles presented in figure 5C and supplementary figure 1B, was determined by manually drawing a line across the max diameter of the vesicle MFI projection image and quantifying the difference between the max intensity of LBR at the vesicle boundary. The size of CD63 vesicle presented in supplementary figure 5C was quantified as Feret diameter of the ROI around individual CD63-positive MVBs, as presented in the zoomed panel of supplementary figure 5A, using Analyze>Measure tool of Fiji image analysis software. The size of CD63 punctae as presented in supplementary figure 5E were quantified using Analyze regions 3D option of MorphLibJ plugin in Fiji software, from the CD63 images thresholded using Max entropy filter. The scale on the expansion microscopy images was adjusted 4X, and the size of LBR positive vesicles was determined by manually drawing an ROI and using the Measure option from the Fiji toolbar to determine the Feret diameter. The 5-LO channel was extracted, intensity thresholded using the Max entropy algorithm, and both the number and mean radii of 5-LO positive objects within the LBR-positive ROI were determined using Analyze>Analyze region 3D option in the MorphLibJ plugin of Fiji software. The data were exported to Graphpad prism and was plotted. The band intensities on the western blots were quantified using the Gels plugin from Analyze toolbar and were plotted in GraphPad prism. To quantify the integrated pixel density of the spots obtained from the dot blots, circular ROI of uniform size was manually created, and the intensity was quantified using Analyze>Measure tool in Fiji image analysis tool. The 3D reconstruction of the z-stack images as presented in Figure 3A and 3C, were performed by exporting the Spectrum LUT (FLAP intensity) and olive pseudo color (Hoechst) images for figure A, and unmodified images for Figure 3C directly to Arivis Vision 4D 3.5 platform. In the software the volumetric rendering of the 3D images were performed using maximum intensity enabling projection of data along view direction. Rendered volumes were tilted to an angle of 125º and rotated along the Y-axis at 25 frames per second containing a total of 125 frames, to create a stereoscopic view.

### Statistics and reproducibility

All the data presented are from at least three independent biological replicates. An appropriate test of significance has been used to determine the level of confidence and variability in the data and mentioned in the corresponding figure legends. All the raw data presented have been provided as “statistics source data” for the respective figures. All the raw images for microscopy data are available from the corresponding author upon reasonable request.

## SUPPLEMENTAL INFORMATION

**Supplementary Figure 1.**
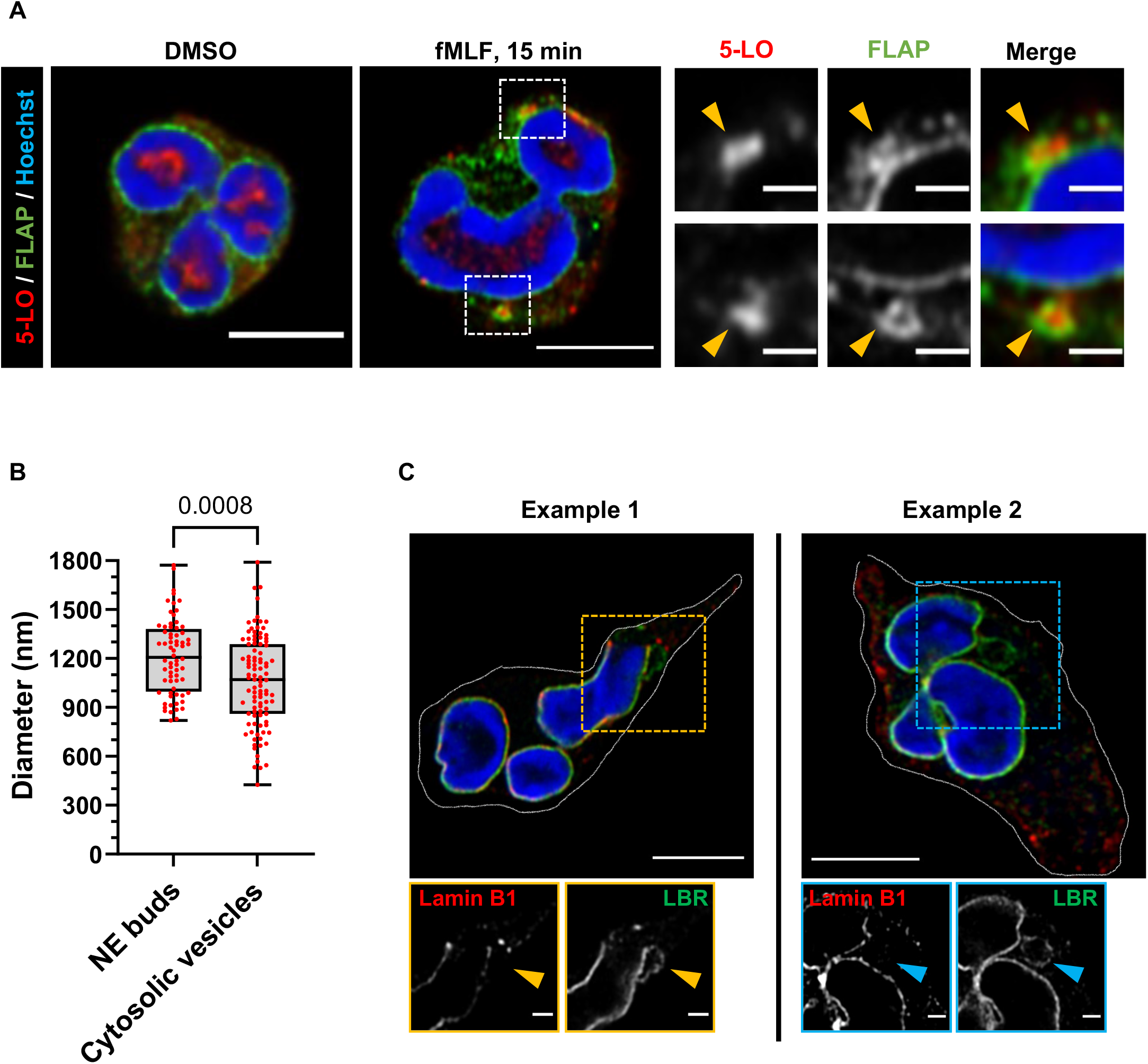
Characterization of LBR-positive NE buds and cytosolic vesicles. **A** Representative Airyscan microscopy image of fixed PMNs uniformly stimulated with either DMSO or 20 nM fMLF showing the distribution of FLAP (green) and 5-LO (red) (n=3). The yellow arrowheads point to nuclear buds. Scale bar is 5 µm, in the inset scale bar is 2 µm. **B** Box-whisker plots showing the size distribution of LBR-positive NE buds and cytosolic vesicles in PMNs chemotaxing towards fMLF (n=5), where each red dot represents the value from the individual bud (68) and cytosolic vesicle (99), plotted as median with range. The indicated P value was determined using the Mann-Whitney test. **C** Examples of fixed PMNs chemotaxing towards 100 nM fMLF and stained for LBR (green), Lamin B1 (red), and Hoechst (blue) acquired using Airyscan microscopy (n=3). Scale bar is 5 µm, in the inset it is 1 µm. NE buds are shown in yellow boxes and cytosolic vesicles are shown in blue boxes. The yellow arrowheads point to nuclear buds and the blue arrowheads point to cytosolic vesicles

**Supplementary Figure 2.**
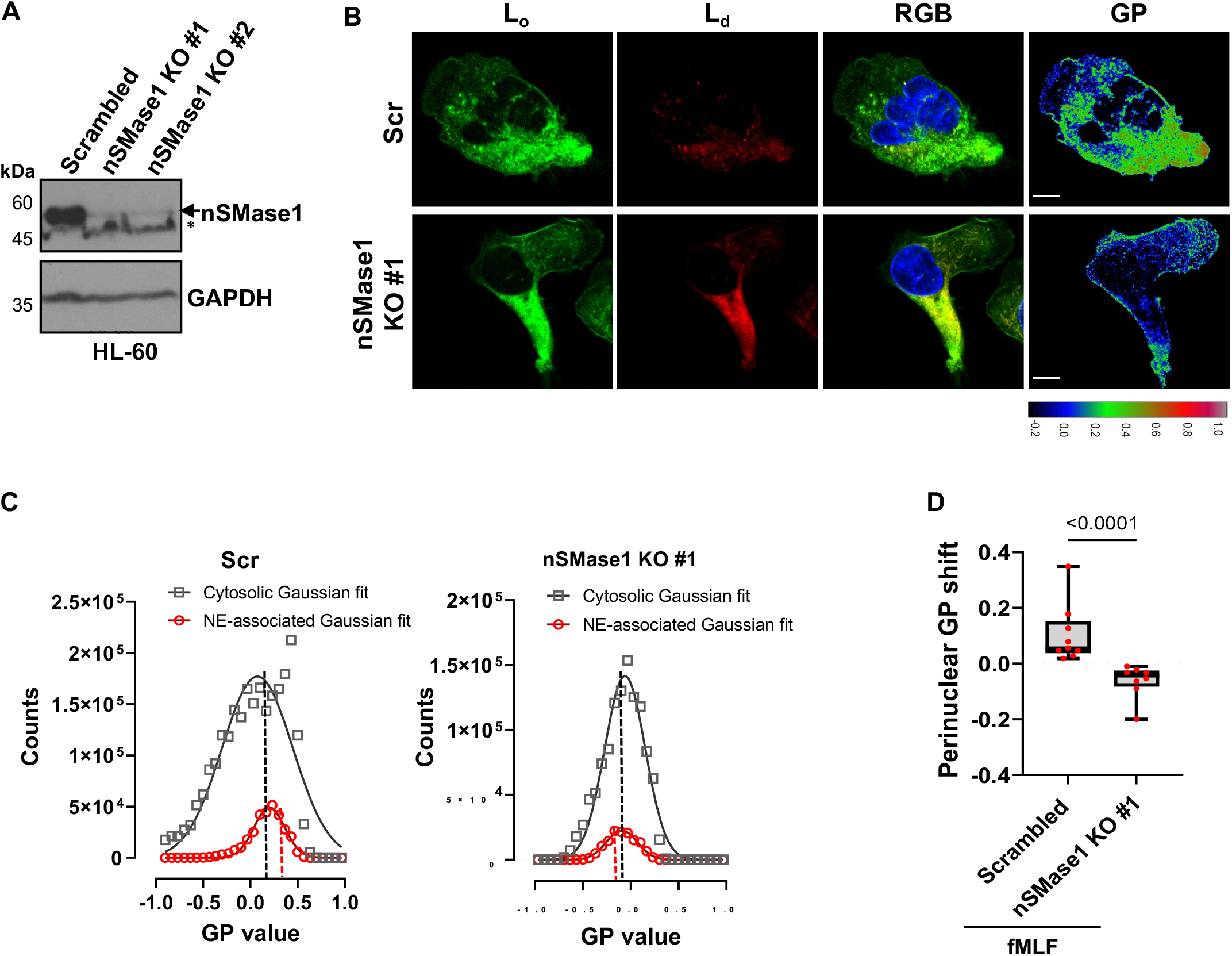
nSMase1 regulates fMLF-induced perinuclear lipid ordering. **A** Representative western blot image showing the levels of nSMase1 in Scr or nSMase1 CRISPR knockout HL-60 cell lysates. GAPDH was used as a loading control (n=2). * Denotes non-specific band detected by the anti-nSMase1 antibody. **B** Representative fluorescence microscopy images, showing the lipid-ordered (L_o_) domains, lipid-disordered (L_d_) domains, RGB images of L_d_, L_o,_ and Hoechst merged, and GP images of either Scr or nSMase 1 KO dHL-60 cells chemotaxing towards 100 nM fMLF stained with di-4ANEPPDHQ (n=3). **C** Graphs depicting the gaussian distribution of cytosolic and perinuclear GP pixel intensity in Scr or nSMase KO dHL-60 cells, as shown in panel B. **D** Box-whisker plots showing the median value of the perinuclear GP intensity distribution obtained from the GP images of Scr and nSMase1 KO dHL-60 cells. Data is quantified from 9 cells out of three independent experiments and is presented as median with range, where each red dot represents the value from one cell. Indicated P value is determined using the Mann-Whitney test.

**Supplementary Figure 3.**
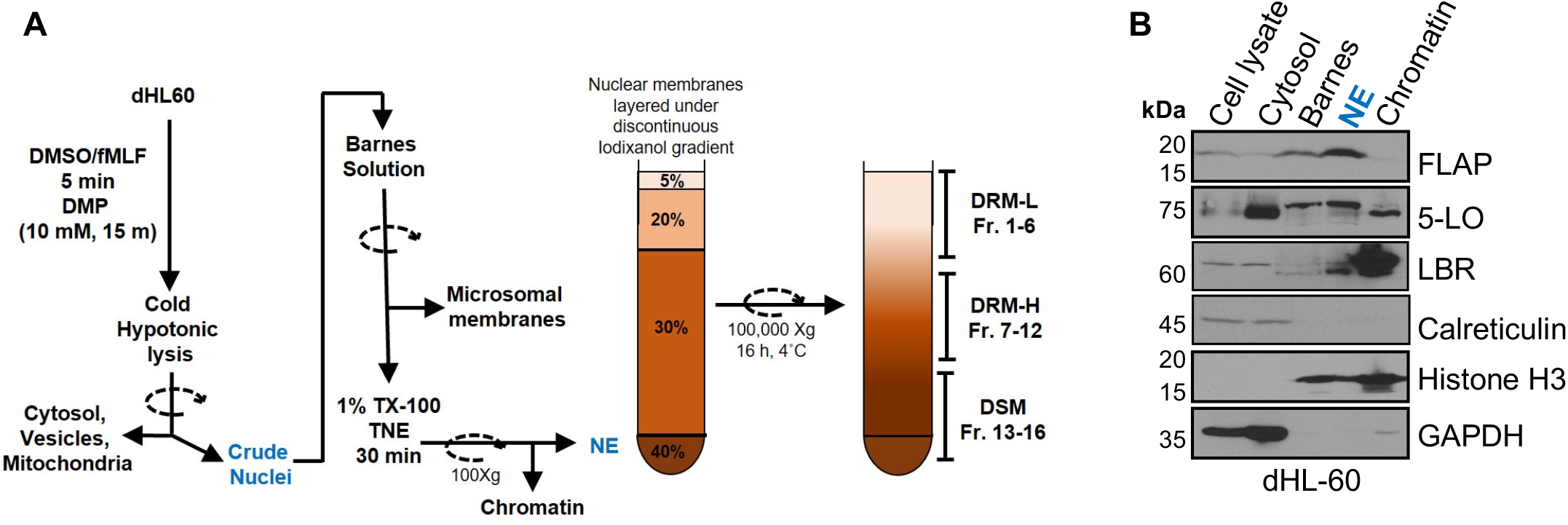
Characterization of NE membranes from activated WT dHL-60 cells. **A** Schematic illustration of the methodology used to isolate DRM and DSM fractions from the isolated NE. **B** Representative western blot showing the efficiency of the fractionation protocol (n=2).

**Supplementary Figure 4.**
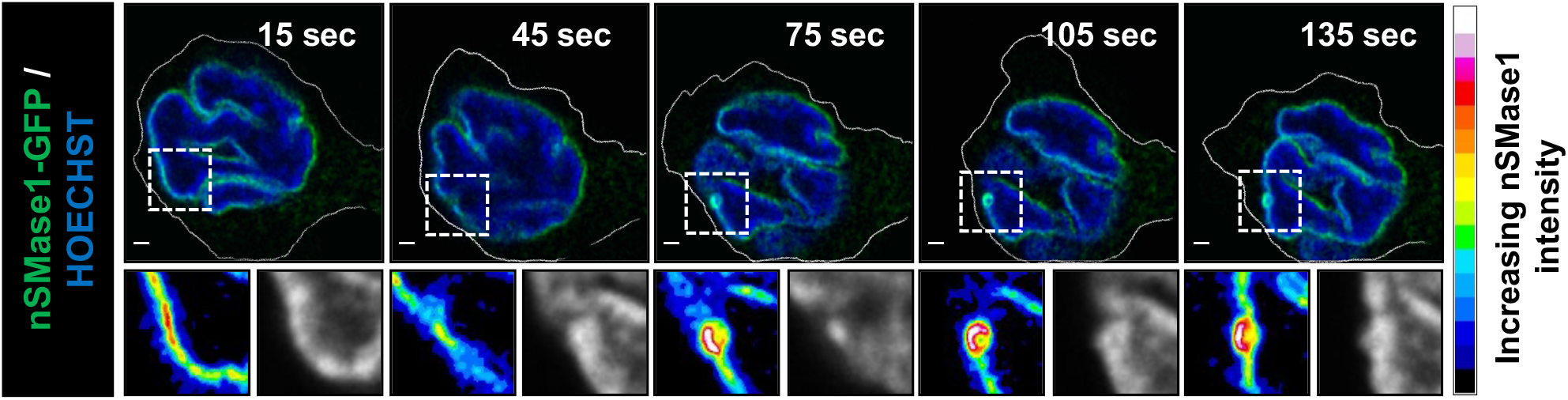
nSMase1-GFP is enriched at sites of nuclear budding. Representative time-lapse images of nSMase1-GFP expressing dHL-60 cells chemotaxing towards 100 nM fMLF. The zoomed section of the images shows the nSMase1-GFP signal as fluorescence intensity spectrum (scale on right) and Hoechst in grayscale. Scale bar is 2 µm. N= 6 cells each acquired from two independent experiments. Also, see Movie S4.

**Supplementary Figure 5.**
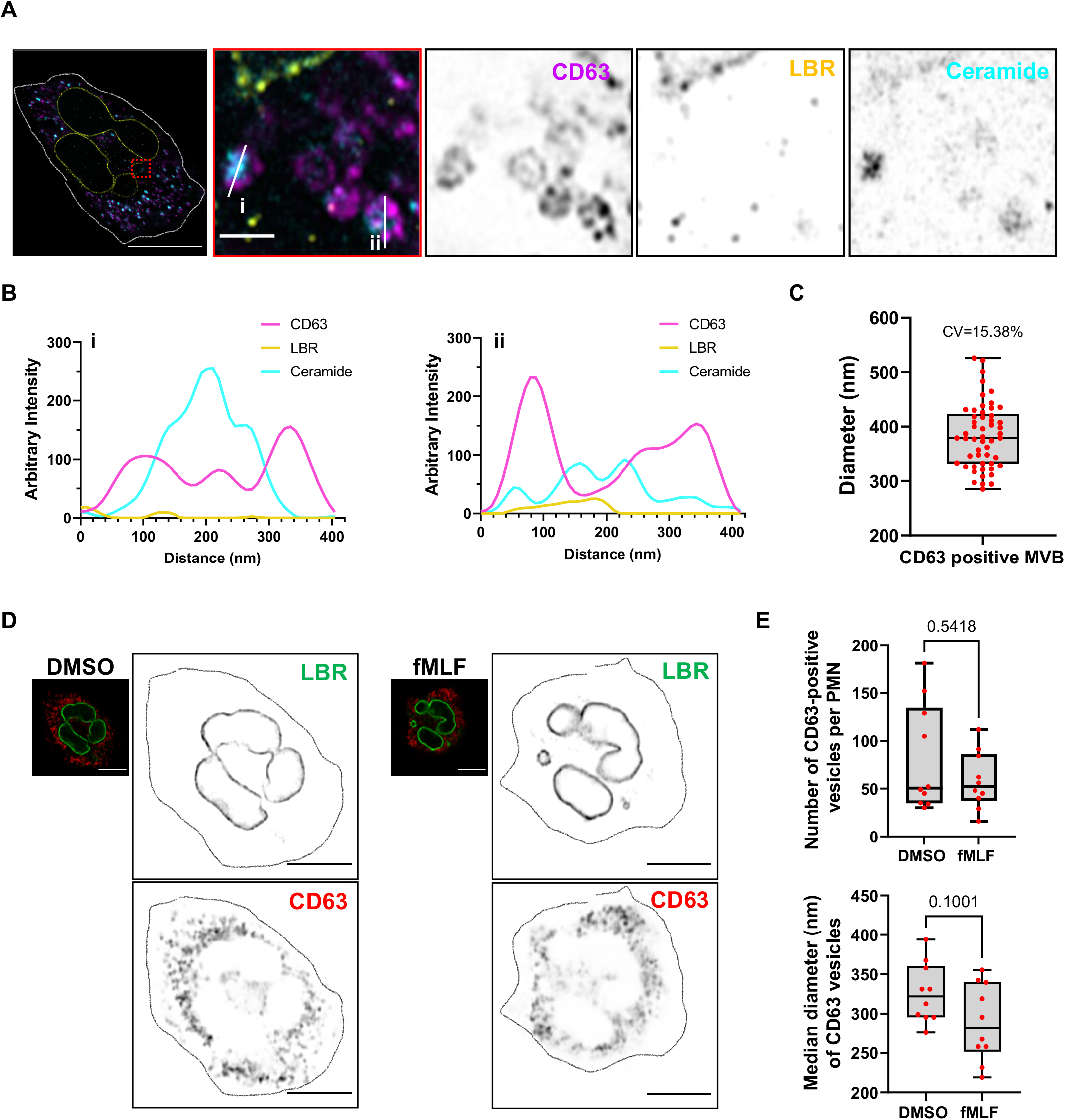
LBR-positive NE buds do not contain CD63. **A** Four-fold expansion microscopy image of fixed human PMNs chemotaxing towards 100 nM fMLF, captured using Airyscan microccopy, stained for CD63 (magenta), LBR (yellow) and ceramide (cyan) (n=3). Scale bar 5 µm. In the inset, the scale bar is 400 nm. **B** Line profiles showing the presence of 50-100 nm ceramide (cyan) positive punctae within the CD63 (magenta) positive conventional MVBs. **C** Box-whisker plot showing the distribution of the median diameter of CD63-positive MVBs. Data are plotted as median with range of 52 CD63-positive MVBs (red dots) from 8 cells pooled from two independent experiments. **D** Representative Airyscan microscopy images of fixed PMNs uniformly stimulated with 100 nM fMLF for 30 min and stained with LBR (green) and CD63 (red) (n=3). Enlarged images are depicted in inverted grayscale. Scale bar is 5 µm. **E** Box-whisker plots showing the number and median diameter of CD63-positive vesicles per cell, as shown in panel C. Data are plotted as median with range and P value calculated using Mann-Whitney test yield non-significant values (n=3).

**Supplementary Figure 6.**
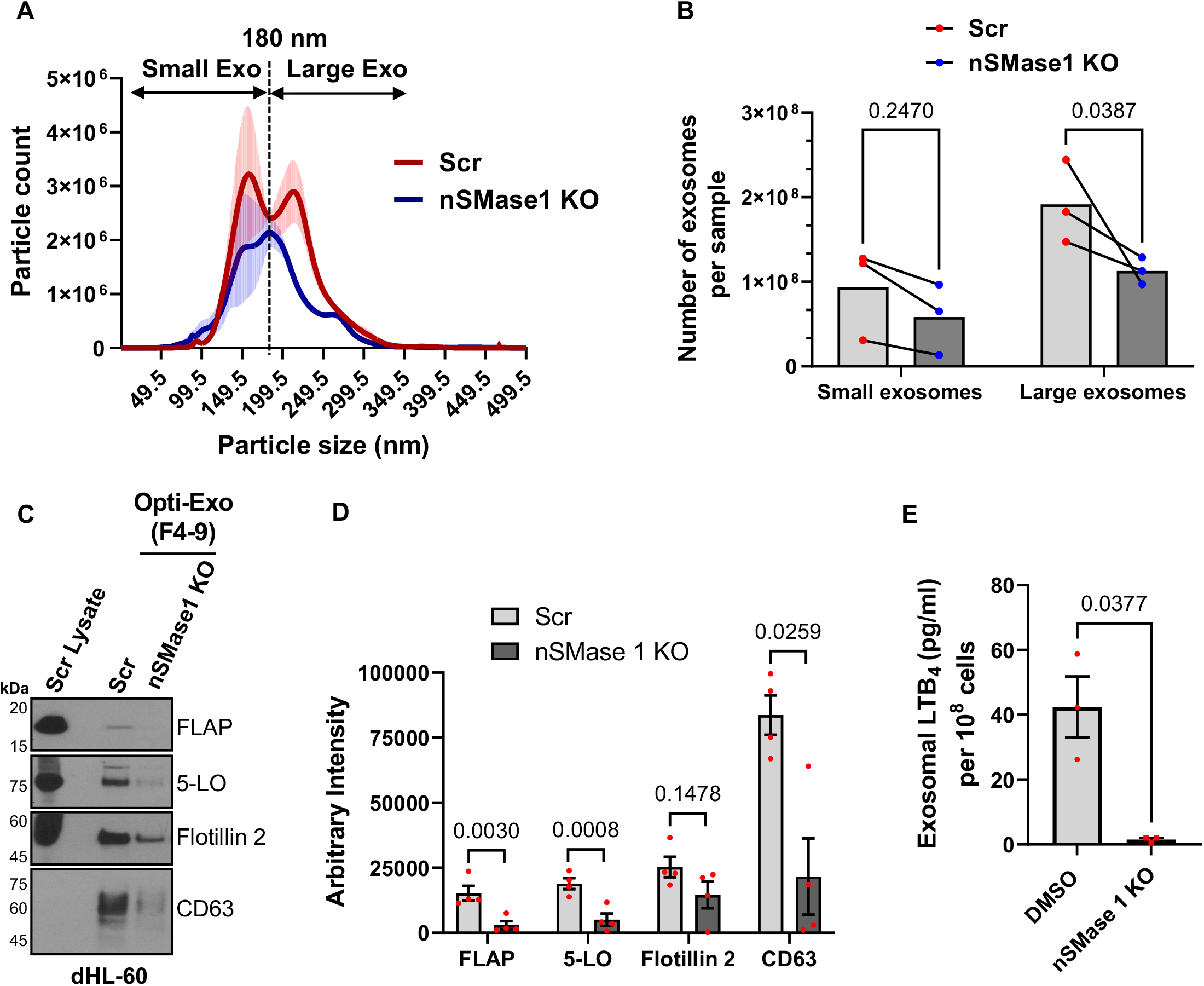
Characterization of exosomes isolated from activated Scr and nSMase1 KO dHL-60 cells. **A** Histogram showing the particle count and size of exosomes purified from either Scr or nSMase1 KO dHL60 cells stimulated with 100 nM fMLF for 15 min. Data were obtained from nanoparticle tracking analysis (NTA) of the isolated exosomes and is plotted from three independent experiments as mean ± SEM. The dotted lines parallel to the y-axis, at 180 nm, indicate the segregation of two exosome populations, small (0-180 nm) and large (181-360 nm). **B** Quantification of the area under the curve from the NTA data. Data from three independent experiments are presented as paired experiments. P-value was obtained two-way RM ANOVA **C** Representative western blot images showing the levels of FLAP, 5-LO, Flotillin 2, and CD63 in pooled fractions 4-9 of density-gradient purified exosomes isolated from either Scr or nSMase1 KO dHL-60 cells stimulated with 100 nM fMLF for 15 min. Scr cell lysate represents the amount of protein from 1/100^th^ the number of cells used for exosome isolation. **D** Bar graph showing the quantifications of the band intensity of FLAP, 5-LO, Flotillin 2, and CD63 in Scr and nSMase1 KO exosomes. Four data points are plotted as mean ± SEM where each red dot represents the value from one experiment. P values determined using ratio paired t-test are reported. **E** Bar graph showing exosomal LTB_4_ levels from the Scr or nSMase1 KO dHL60 cells stimulated with 100 nM fMLF for 15 min. Data from three independent experiments are plotted as mean ± SEM. P value was obtained using ratio paired t-test.

**Supplementary Figure 7.**
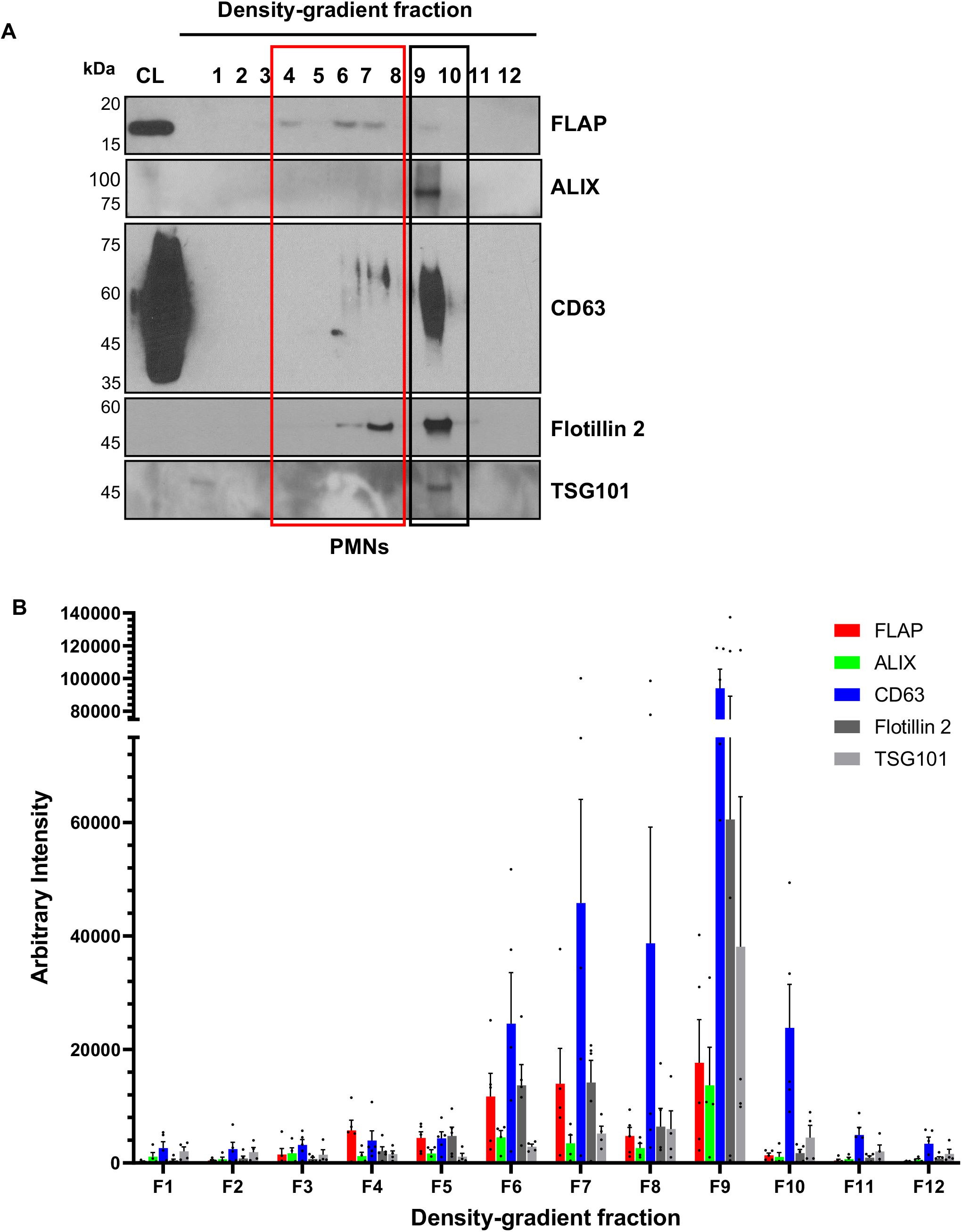
Characterization of exosomes isolated form activated PMNs. **A** Representative western blot showing the distribution of FLAP, CD63, Flotillin 2, TSG101, and ALIX in various fractions of density-gradient purified exosomes isolated from the supernatant of PMNs stimulated with 100 nM fMLF for 15 min (n=4-5). **B** Bar graph showing the arbitrary band intensity of the indicated proteins. Data are plotted as mean ± SEM of at least three independent experiments.

**Supplementary Figure 8.**
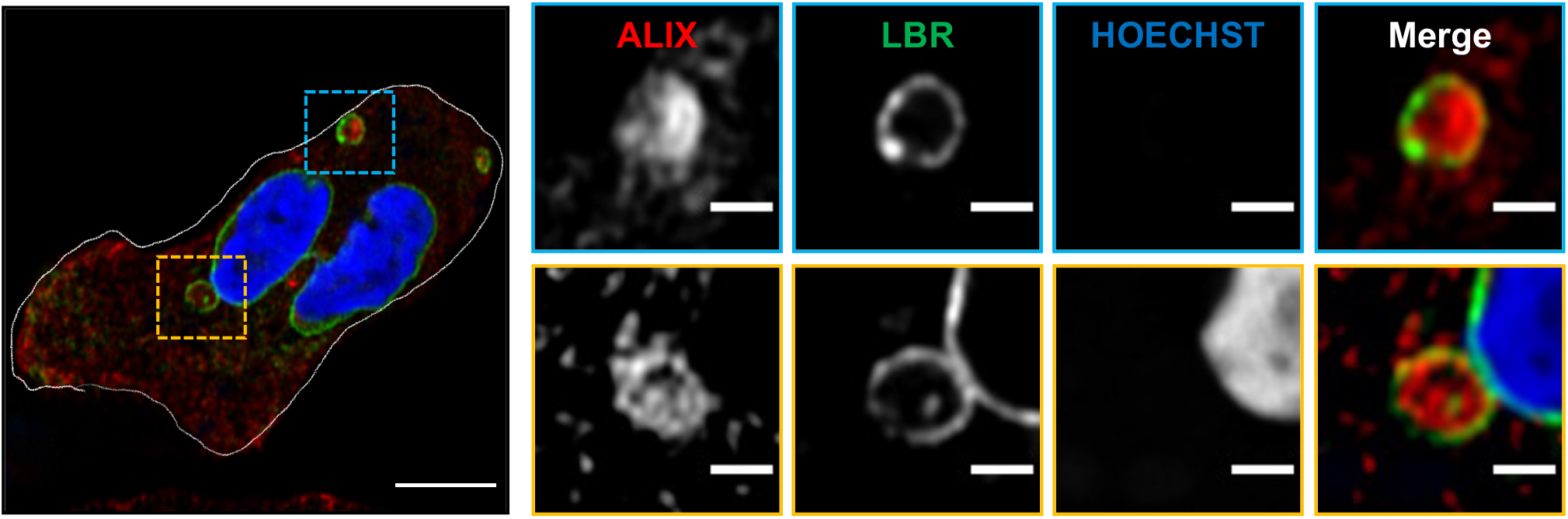
ALIX and LBR distribution in activated PMNs. Representative Airyscan microscopy images of fixed PMNs chemotaxing towards 100 nM fMLF and stained for LBR (green) and ALIX (red) (n=3). Scale bar is 5 µm. In the inset, the scale bar is 1 µm. NE buds are shown in yellow boxes and cytosolic vesicles are shown in blue boxes.

**Movie S1**

Stitched time-lapse images of di-4ANEPPDHQ stained PMNs chemotaxing towards 100 nM fMLF, showing L_o_ regions (green, left), L_d_ regions (red, center), and the corresponding GP image. Pseudo-colored GP images are based on the colormap, with white/red shades depicting lipid ordered regions and blue/darker shades representing lipid disordered regions. For colormap, see Figure 2A. Images were captured at 15-sec intervals and stitched together to create a movie at 20 frames per sec speed.

**Movie S2**

Stereoscopic rendering of olive-colored Hoechst-stained isolated nuclei, showing the intensity distribution of FLAP staining, as presented in Figure 3A.

**Movie S3**

Stereoscopic rendering of FLAP, ceramide, and Hoechst 33342 stained isolated nuclei, showing the differential localization of FLAP and ceramide, as presented in Figure 3C.

**Movie S4**

Time-lapse movie of dHL60 cells expressing nSMase1-eGFP (green) and stained with Hoechst (blue) chemotaxing towards 100 nM fMLF. Images were captured at 15-sec intervals and stitched together at frame rate of 2 frames per sec.

